# Multi-level Bayesian analysis of monk parakeet contact calls shows dialects between European cities

**DOI:** 10.1101/2022.10.12.511863

**Authors:** Simeon Q. Smeele, Stephen A. Tyndel, Lucy M. Aplin, Mary Brooke McElreath

**Author notes:** shared first author (Simeon Q. Smeele, Stephen A. Tyndel). shared senior author (Lucy M. Aplin, Mary Brooke McElreath).

## Abstract

Geographic differences in vocalisations provide strong evidence for animal culture, with patterns likely arising from generations of social learning and transmission. The current knowledge on the evolution of vocal variation has predominantly focused on fixed repertoire, territorial song in passerine birds. The study of vocal communication in open-ended learners and in contexts where vocalisations serve other functions is therefore necessary for a more comprehensive understanding of vocal dialect evolution. Parrots are open-ended vocal production learners that use vocalisations for social contact and coordination. Geographic variation in parrot vocalisations typically take the form of either distinct regional variations known as dialects or graded variation based on geographic distance known as clinal variation. In this study, we recorded monk parakeets (*Myiopsitta monachus*) across multiple spatial scales (i.e. parks and cities) in their European invasive range. We then compared calls using a multi-level Bayesian model and sensitivity analysis, with this novel approach allowing us to explicitly compare vocalisations at multiple spatial scales. We found support for founder effects and/or cultural drift at the city level, consistent with passive cultural processes leading to large scale dialect differences. We did not find a strong signal for dialect or clinal differences between parks within cities, suggesting that birds did not actively converge on a group level signal, as expected under the group membership hypothesis. We demonstrate the robustness of our findings and offer an explanation that unifies the results of prior monk parakeet vocalisation studies.

## Introduction

Differences in vocalisations between groups or populations have been identified across multiple animal species. Such geographic variation in vocalisations has provided some of the strongest evidence for vocal learning and animal culture (Marler and Tamura 1962; Catchpole and Slater 2003; Podos and Warren 2007; Aplin 2019). In particular, patterns of vocal variation in songbirds have been the focus of decades of intensive research (Slater 2003). In songbirds, song is primarily used to defend territories and attract mates (Krebs and Kroodsma 1980; Kroodsma and Byers 1991; Catchpole and Slater 2003), and is often exclusively learned early in development. Coupled with vocal convergence and conformity (Lachlan, Ratmann, and Nowicki 2018), this early flexibility can result in highly stable and localised dialects. For example, male new world sparrows (*Passerllidae*) produce complex songs that form clear geographic dialects (Williams, Levin, et al. 2013; Lachlan, Ratmann, and Nowicki 2018). These dialects are maintained over long periods of time and may play an important function in species recognition and mate choice (Lachlan, Ratmann, and Nowicki 2018; Slater 2003). Furthermore, the way dialects are structured can depend heavily on behavior and social structure. This is supported by examples of species that have limited migration and dispersal between populations, which show a gradual change in vocal differentiation across a geographical clinal gradient (D. E. Irwin, Thimgan, and J. H. Irwin 2008). However, the study of vocal variation in open-ended vocal production learners outside the context of bird-song is relatively understudied and the mechanisms leading to emergent dialect or clinal patterns in these cases are poorly understood (Wright and Dahlin 2018).

Open-ended vocal production learning refers to the ability to modify or change produced vocalisations throughout adulthood (Beecher and Brenowitz 2005; Janik and Slater 1997; Janik and Knörnschild 2021). Open-ended vocal production learning has evolved in several taxonomic groups including bats, cetaceans and three main groups of birds: hummingbirds (*Trochilidae*), passerines (i.e., *Corvidae, Fringillidae, Sturnidae*) and parrots (*Psittaformes*). Many parrot species show geographic variation in their contact calls (Wright 1996; Wright and Dahlin 2018), and, in captive studies, are able to actively converge their vocalisations across long (multiple weeks) time scales (Hile, Plummer, and Striedter 2000). This observation of group convergence has been hypothesized to lead to group-level vocal signatures (Dahlin et al. 2014). In addition to long time scales, parrots can also rapidly modify their calls (i.e, within seconds) (Balsby and Bradbury 2009; Thomsen, Balsby, and Dabelsteen 2019; Scarl and Bradbury 2009; Vehrencamp et al. 2003) depending on specific social context (for example, addressing flock members (Balsby, Momberg, and Dabelsteen 2012)). This extreme rapid flexibility could be another possible mechanism leading to overarching geographic variation (Barker et al. 2021).

Several hypotheses have been proposed to explain patterns of geographic vocal variation in openended vocal production learners such as parrots. The *group membership hypothesis* posits that vocal dialects serve a functional purpose of increased recognition of group members and possibly foraging efficiency within social groups (Bradbury, Vehrencamp, et al. 1998; Sewall, Young, and Wright 2016). In support of this hypothesis, a wide range of studies have found that some parrot species (Wright and Dorin 2001; Vehrencamp et al. 2003; Dahlin et al. 2014), bats (Knörnschild et al. 2012) and dolphins (Janik and Slater 1998) appear to use calls to strengthen social bonds in groups. Under this framework, particular call types, and/or dialects could undergo social selection, allowing for stable call types (Wright 1996). In terms of observable predictions, we would propose that this active process of group convergence should manifest as group signatures at small geographic scales, with this scale further depending on group size and social structure. Along the same lines, if populations demonstrate large degrees of fission-fusion dynamics, this could possibly lead to a clinal gradient, where vocalisations produced in close geographic proximity sound more similar than those produced further apart (Bradbury, Cortopassi, et al. 2001).

The *cultural drift hypothesis* proposes that vocal variation forms as the result of passive cultural processes, with either copying errors or innovations combined with neutral or directional cultural evolution that allows for groups to diverge (Williams, Levin, et al. 2013; Williams and Lachlan 2022; Payne 1978). Previous research suggests that sexual (Nowicki, Peters, and Podos 1998) and social selection (Lachlan, Ratmann, and Nowicki 2018) both represent likely selective pressures in songbird species. In open-ended learning species such as parrots, contact calls are likely not subject to sexual selection (Bradbury and Balsby 2016). Isolation and cultural drift combined with social selection, therefore appears to be the most plausible explanation for many species. For example, crimson rosella (*Platycercus elegans*) (Ribot et al. 2012) and St. Lucia parrots (*Amazona versicolor*) (Martínez and Logue 2020) both demonstrate dialect boundaries that correspond with barriers to movement. Unlike the *group membership hypothesis*, the *cultural drift hypothesis* does not necessarily require selection for convergence at the group level. Under this scenario, we would expect to observe dialects across isolated geographic regions, likely at larger scales where boundaries exist that isolate populations.

In contrast to the *group membership hypothesis*, the *individual signature hypothesis* posits that individuals actively modify their vocalisations to try and sound as distinct from one another as possible (Nowicki and Searcy 2014). In this scenario, we would not necessarily expect to observe geographic vocal variation, despite the social learning of vocalisations. This is because the drive for individual distinctiveness may lead to fully maximised variation within groups (Gillam and Chaverri 2012), making any effect of cultural drift between populations difficult to detect. This type of pattern has been observed in other open-ended learning species such as dolphins (Oswald et al. 2021), and parrot species such as green rumped parrotlets (*Forpus passerinus*) (Berg, Delgado, Okawa, et al. 2011) and monk parakeets (Smith-Vidaurre, Araya-Salas, and Wright 2020). However, the *individual signature hypothesis* is not necessarily mutually exclusive with the *group membership hypothesis*. Indeed there is evidence that some species can maintain individual signatures while also maintaining strong group level signatures (Wright 1996; Thomsen, Balsby, and Dabelsteen 2013; Wright 1996). The precise mechanism that causes individual signatures to outweigh dialects versus having strong individual signatures in concert with strong dialect boundaries remains unclear.

Monk parakeets (*Myiopsitta monachus*) are an excellent study system to elucidate the processes that lead to geographic vocal variation in open-ended vocal learners. Monk parakeets have a large invasive range across Europe and North America (Forshaw and Cooper 1989), where populations are largely concentrated in cities, often with little movement between them (Postigo et al. 2019; Edelaar et al. 2015). Importantly, monk parakeets have various population substructures that allow for close study of geographic vocal variation patterns at multiple scales. They nest in single or compound nests, the latter containing multiple nests, each with one or multiple chambers per pair (SQS personal observation). Nest openings correspond to nest chambers, which can serve as a proxy for population size. These nest structures occur in larger nesting colonies. The term colony is often defined as one or more nest structures located within 200m of each other (see (Reed et al. 2014)). In cities and invasive populations, these nesting colonies are often located within parks or other green areas, clearly delineated from other colonies (Eberhard 1998), although with potential between-park movement and dispersal (Bucher, Martin, et al. 1990). A recent study in the native range of monk parakeets found evidence that individual signatures outweighed any emergent dialects (Smith-Vidaurre, Araya-Salas, and Wright 2020). Interestingly, regional dialects between cities have been observed in the invasive populations of monk parakeets in the United States (Buhrman-Deever, Rappaport, and Bradbury 2007).

In the current study, we aim to assess these competing hypotheses by examining patterns of vocalisations across parks and cities in the invasive range of monk parakeets in Europe. Because most European populations of monk parakeets have comparable genetic compositions (Edelaar et al. 2015), it allows us to consider the influence of cultural processes rather than potential genetic differences between the populations. Our populations contain many sub-populations (i.e., parks) making it possible to conduct a two-level comparison with many replicates. If dialects or clinal variation are found at the park level, selection for call sharing with other group members is likely at play, lending credence to vocal convergence via the *group membership hypothesis*. Of course, if movement between parks is low, one could not rule out the possibility of cultural drift also occurring.

If dialects exist only at the city level, it would suggest a cultural founder effect and/or cultural drift, similar to that often observed in songbirds (Lachlan, Ratmann, and Nowicki 2018; A. J. Baker and Jenkins 1987). Lastly, if no dialects are found between cities, several testable hypotheses, such as founder effects and selection for distinctive calls at the individual level (Smith-Vidaurre, Araya-Salas, and Wright 2020) could be considered.

## Methods

### Study System

Monk parakeets are a medium sized colonially-nesting parrot. While native to South America, they have been transported by the pet trade across the world and have established large invasive populations in several European countries including Spain, France, Belgium, Italy and Greece. These populations are usually clustered in cities and towns, often with relatively little dispersal between them (Dawson Pell et al. 2021). Monk parakeets in Europe breed from March to August and roost in their nests year-round (Senar, Carrillo-Ortiz, et al. 2019). Nests are often highly spatially clustered, with several nest chambers per nest, several nests per tree and trees often clustered together (Eberhard 1998). Population sizes vary within and between cities and parks, with estimates ranging between one nest chamber in Thisio park, Athens to 99 in Gendarmerie School Park, Athens.

### Data Collection

We collected vocalisations from monk parakeets in 39 parks across eight cities in four countries: Athens, Barcelona, Bergamo, Brussels, Legnago, Madrid, Pavia and Verona in November 2019 (see Table 1 for sampling effort per park, see Figure 1 for sampling area, and see supplemental materials for maps of parks within cities). Vocalisations were opportunistically recorded between sunrise and sunset with a Sennheiser K6 + ME67 microphone and either a Sony PCM M10 or Sony PCM D100 recorder. Recordings were made at a distance between one and 20 meters and lasted 20 minutes or until the bird moved away. If calls could be assigned with certainty to a focal bird this was verbally annotated.

**Table 1:** Recording effort per city and park. Number of days represents how many days the parks were visited. Not all recording sessions were entire days. Number of calls represents how many calls were included in the final analysis.

| city | park | number of days | number of calls | number of nest openings |
| --- | --- | --- | --- | --- |
| Athens | Oluf Palme Playground | 3 | 35 | 9 |
| Athens | National Garden | 4 | 287 | 49 |
| Athens | Alsos Ilision | 2 | 52 | 10 |
| Athens | Gendarmerie School Park | 3 | 86 | 99 |
| Athens | Thissio Park | 1 | 2 | 1 |
| Barcelona | Parc de la Ciutadella | 3 | 85 | 33 |
| Barcelona | Jardins del Turo del Putxet | 1 | 98 | 1 |
| Barcelona | Jardins de Ghandi | 1 | 2 | 4 |
| Barcelona | Jardins de Josep Trueta | 1 | 20 | 7 |
| Barcelona | Parc Grande de Sant Marti | 1 | 44 | 54 |
| Barcelona | Jardin de la Maternitat | 1 | 19 | 11 |
| Bergamo | Faunistic Park Le Cornelle | 2 | 456 | 26 |
| Brussels | Parc de Forest | 4 | 559 | 96 |
| Brussels | Ten Reuken | 2 | 19 | NA |
| Brussels | Avenue Louise | 1 | 6 | 7 |
| Brussels | Tenenbosch Park | 1 | 10 | 1 |
| Brussels | Place Guy D'Arezzo | 2 | 107 | 13 |
| Legnago | Legnago | 2 | 345 | 10 |
| Madrid | Parque de el Ritero | 1 | 13 | NA |
| Madrid | Parque de Berlin | 3 | 218 | 65 |
| Madrid | Lago Casa del Campo | 2 | 91 | 18 |
| Madrid | Parque Azorin | 2 | 141 | 55 |
| Madrid | Parque Emperatriz Maria de Austria | 1 | 45 | NA |
| Madrid | Parque Infantil Portalegre | 1 | 9 | 6 |
| Madrid | Quintos de Molinos | 1 | 10 | 2 |
| Madrid | Parque Alfredo Kraus | 1 | 5 | 10 |
| Pavia | Oasi di Sant' Alessio | 1 | 756 | 34 |
| Verona | Parco Natura Viva | 1 | 110 | 37 |

**Figure 1.**
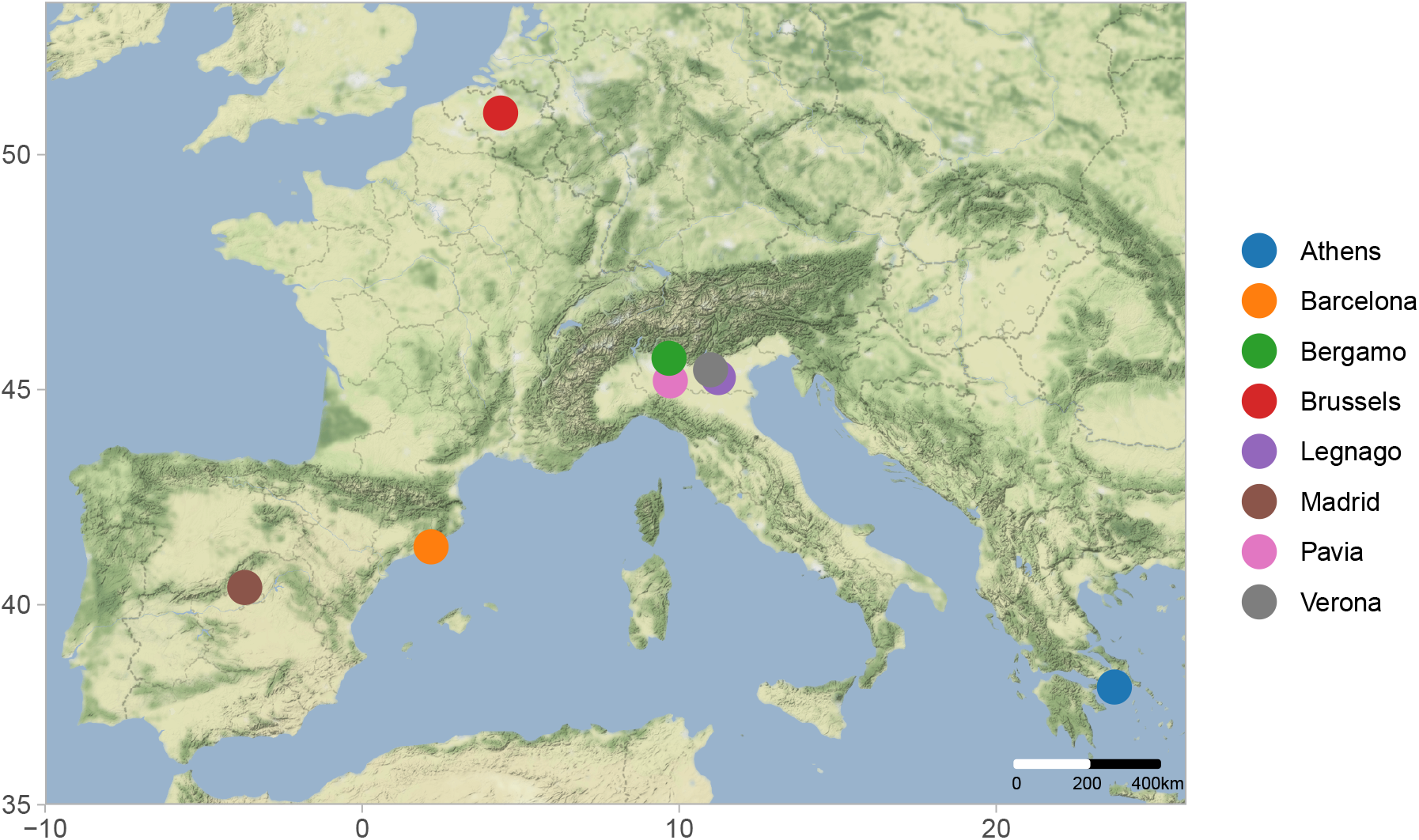
Map of study locations. Map created using ggmap (Kahle and Wickham 2013), ggrepel (Slowikowski 2021), and ggsn (Santos Baquero 2019). Map tiles by Stamen Design, under CC BY 3.0. Data by OpenStreetMap, under ODbL.

Although individuals were not identifiable across recordings, whenever possible we recorded the vocalising individual with a unique ID within a recording. We also included recordings when the vocalising individual was not assigned a unique ID. In order to avoid assigning a unique ID to each vocalisation made by an unidentified individual, we grouped them by five minute intervals of recording, assuming recordings during that time span came from one individual. Some recordings were also videotaped with a Philips HC-V777EG-K to allow assignment of calls during processing. We tested how this incorrect pooling might have affected the results in a sensitivity analysis (see Supplemental Materials).

### Data Processing

Raw recordings were first imported to Raven Lite (Cornell Lab of Ornithology, NY 2016). We manually selected the start and end times of all vocalisations with reasonable signal to noise ratio and annotated the caller ID and behaviour if available. Using a custom script in R (R Core Team 2021), all selected calls from Raven were clipped and high quality spectrograms were created (see Data availability statement). All spectrograms were then manually inspected and calls that were considered to be poor quality were removed.

The remaining calls were categorised as contact calls (tonal calls with at least three peaks in their frequency modulation) and other calls. Contact calls were further manually sorted into six variants: *typical* (stereotyped call with four rounded frequency modulated components), *four triangle* (stereotyped call with four triangular shaped frequency modulated components), *ladder start* (call with low frequency harmonic in the first component), *ladder middle* (call with low frequency harmonic in the middle of the call), *ladder multiple* (call with multiple low frequency harmonic components) and *mix alarm* (call with frequency modulated components mixed with amplitude modulated components). For examples of four of the variants see Figure 2. We choose to use a structural definition to designate call types rather than a behavioural one, since most recordings where behaviours were available were of single perched individuals (Smith-Vidaurre, Araya-Salas, and Wright 2020).

**Figure 2.**
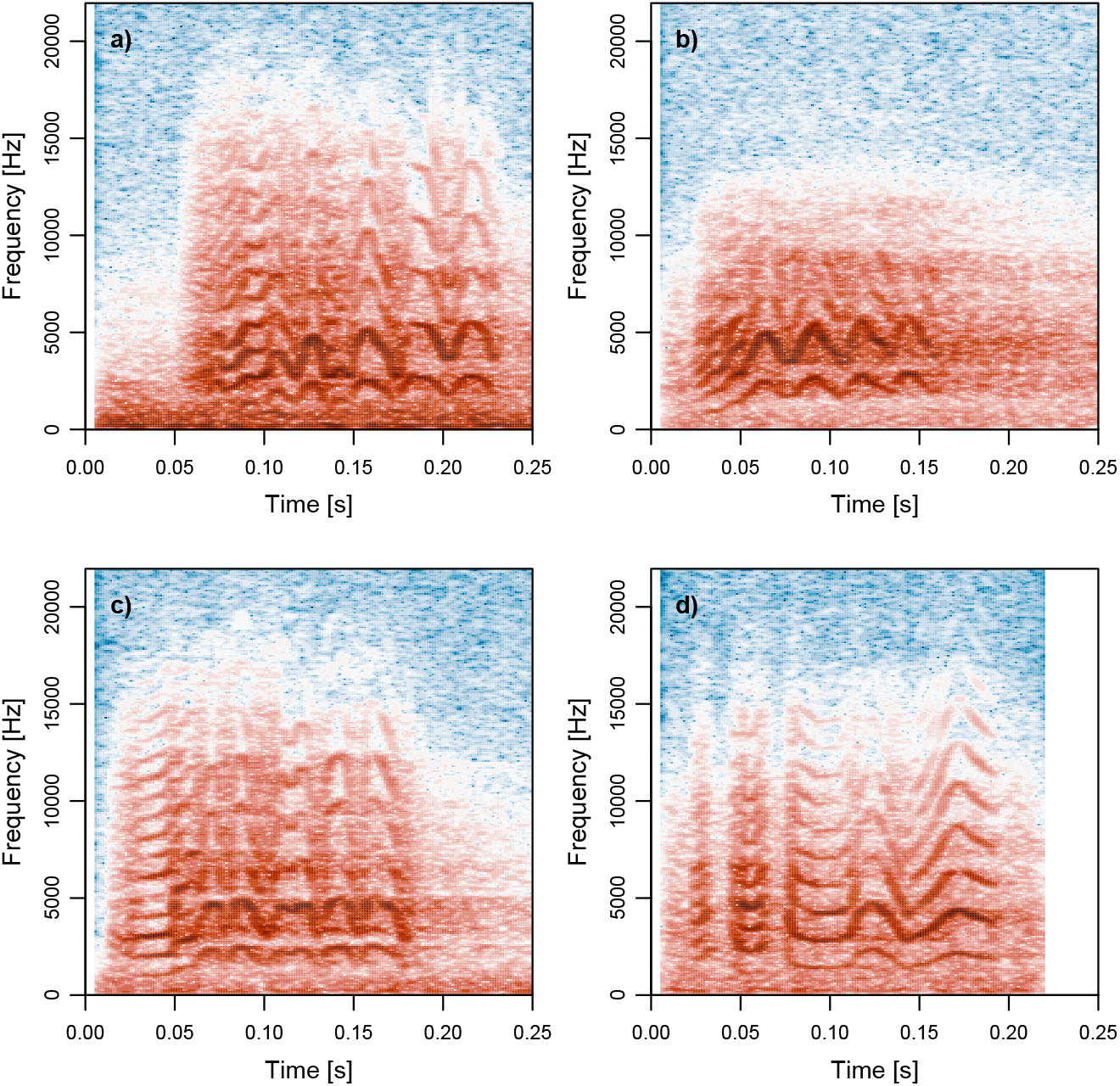
Examples of four contact call variants. a) typical, b) four triangle, c) ladder start, d) mix alarm.

To assess whether our categorizations of call variants were reproducible, we created a randomized sample of 1000 calls from our dataset, including both contact and non-contact calls. We then asked an independent observer to classify the calls. We assessed both how the observer’s classifications of contact calls vs. non-contact calls, and how the observer’s classifications of contact call variants compared to our own. The agreement between our own observations of contact vs. non-contact calls and the independent observers’ observations was very strong (Kappa statistic, k = 0.83, Z = 26.2, %-agree = 91.6). The agreement between our classifications of contact call variants and the independent observers’ classifications was moderately strong (Kappa statistic, k = 0.59, Z = 35.6, %-agree = 74.3).

All good quality contact calls were saved as separate sound files and imported to Luscinia (Lachlan 2007). Using Luscinia’s algorithm we traced the fundamental frequency semi-manually. Some calls could not be traced well and were excluded (28%). The fundamental frequency traces were imported to R and smoothed in two steps to get rid of small errors. First, gaps where Luscinia could not detect the fundamental frequency were filled with a straight line from the last detected point to the first detected point after the gap. Then smooth.spline (*stats*) was used with spar = 0.4 to remove outliers. Traces were visually inspected to ensure proper fit.

We used dynamic time warping (DTW) to measure similarity between all pairs of contact calls. This algorithm takes two time series and measures the optimal similarity between them (Bellman and Kalaba 1959). We used the function *dtw* from the package *dtw* (Giorgino 2009) to run DTW on the fundamental frequency traces. We normalised and log transformed the resulting distance matrix. To represent each call as a single point in two-dimensional space we ran a principal coordinate analysis (PCO) using the function *pcoa* from the package *ape* (Paradis and Schliep 2019). To verify the robustness of our DTW-PCO analysis, we also obtained a distance matrix using spectrographic cross correlation using the entire spectrogram. We used both uniform manifold approximation (UMAP) and principal component analysis (PCA) for dimension reduction (see Supplemental Materials). All approaches gave similar results.

### Statistical Analysis

We used a Bayesian multilevel model to test how much variation in PC1 and PC2 was explained by the two geographic levels of interest, park and city. Both were included as varying effects. To control for pseudo replication we included the verbally annotated IDs whenever possible as varying effects as well. When IDs were not available, we grouped all calls occurring in the same five minute interval as one individual. We conducted a sensitivity analysis to test how well this approach could mitigate the effects of pseudo replication (see Supplemental Materials). The full model structure for PC1 (standardised) is as follows:

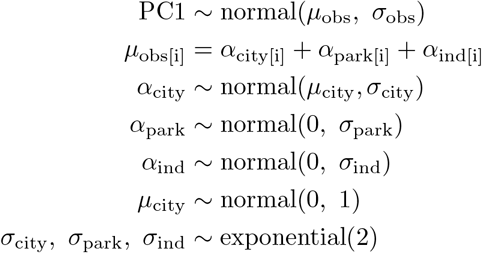

The model was fitted using the No U-turn Sampler, an improved version of the Hamiltonian Monte Carlo algorithm in Stan (Gelman, Lee, and Guo 2015). A similar model was run for PC2.

## Results

In total we traced 3630 contact calls using Luscinia. This encompassed 1-4 days of recording effort and 2-756 recorded calls at each park, with a median of 48.5 calls (n=28 parks). At the city level between 100 (Verona) and 701 calls (Brussels) were recorded, with a median of 459 calls (n=8 cities). See Table 1 for additional sampling details.

There was clustering by city (see Figure 3b) with distinct differences based on PC1 (see Figure 4c). In particular, Bergamo, Legnago and Pavia were different from the other cities (Figure 4c). For the second principal coordinate, the results demonstrated high levels of differentiated dialects between the majority of different cities (see Figure 4a). In general, there was considerable evidence that vocalisations varied between cities (mean *s*_ciy_ PC 1: 0.40, 89% PI: 0.19-0.67, mean *s*_city_ PC 2: 0.58, 89% PI: 0.34-0.92), and varied less between parks (mean *s*_park_ PC 1: 0.21, 89% PI: 0.12-0.34, mean *s*_park_ PC 2: 0.29, 89% PI: 0.19-0.42) as demonstrated by the sigma parameters and pair-wise contrasts (see Figure 3). These results were consistent across methods (see Supplemental Materials).

**Figure 3.**
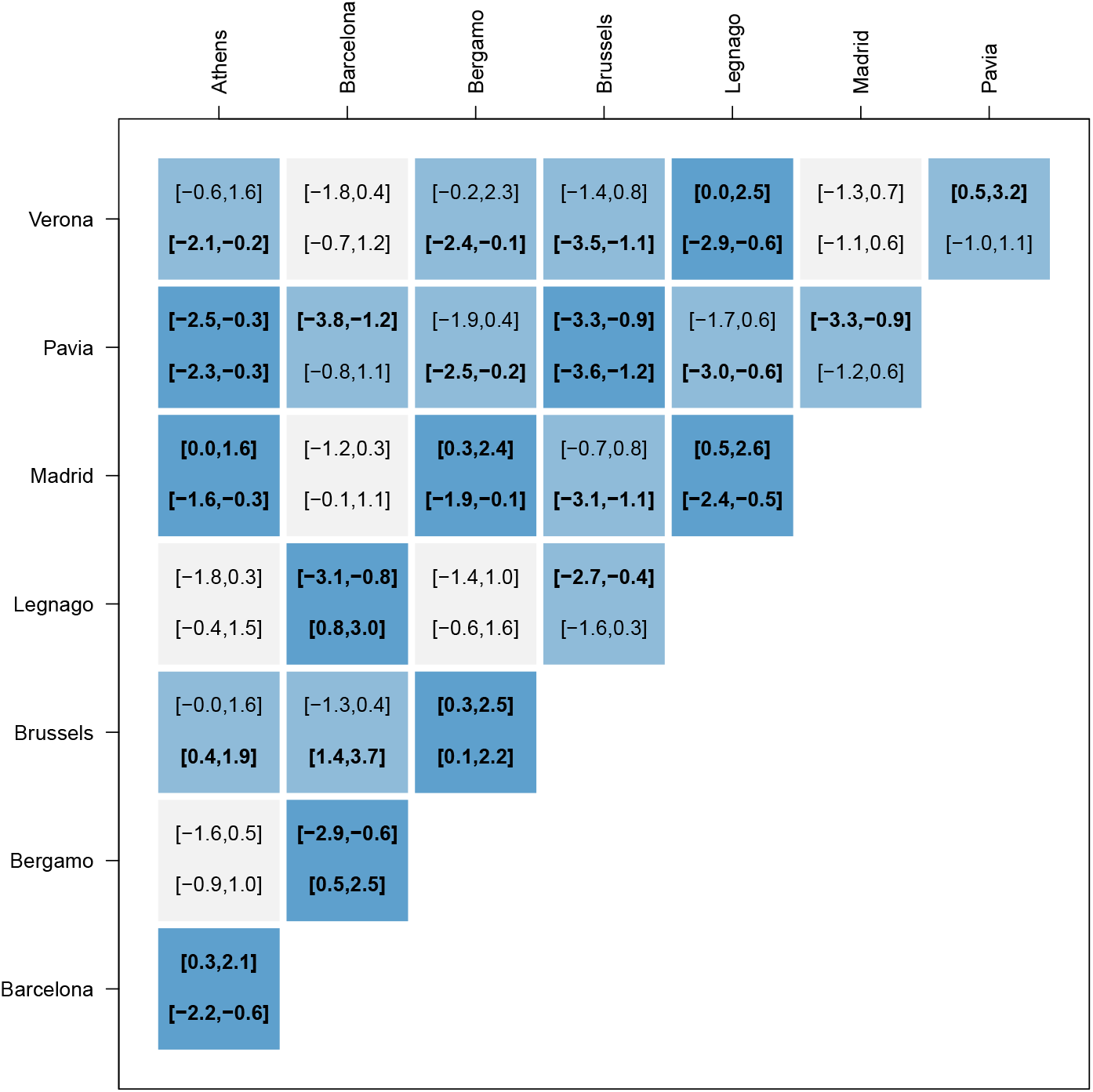
Pairwise contrasts between city means. Number in brackets give the 89% posterior interval for principal coordinate 1 (top) and 2 (bottom) for all city pairs. Intervals are in bold if they do not overlap 0. Squares are coloured dark if either one or both of the intervals do not overlap 0.

**Figure 4.**
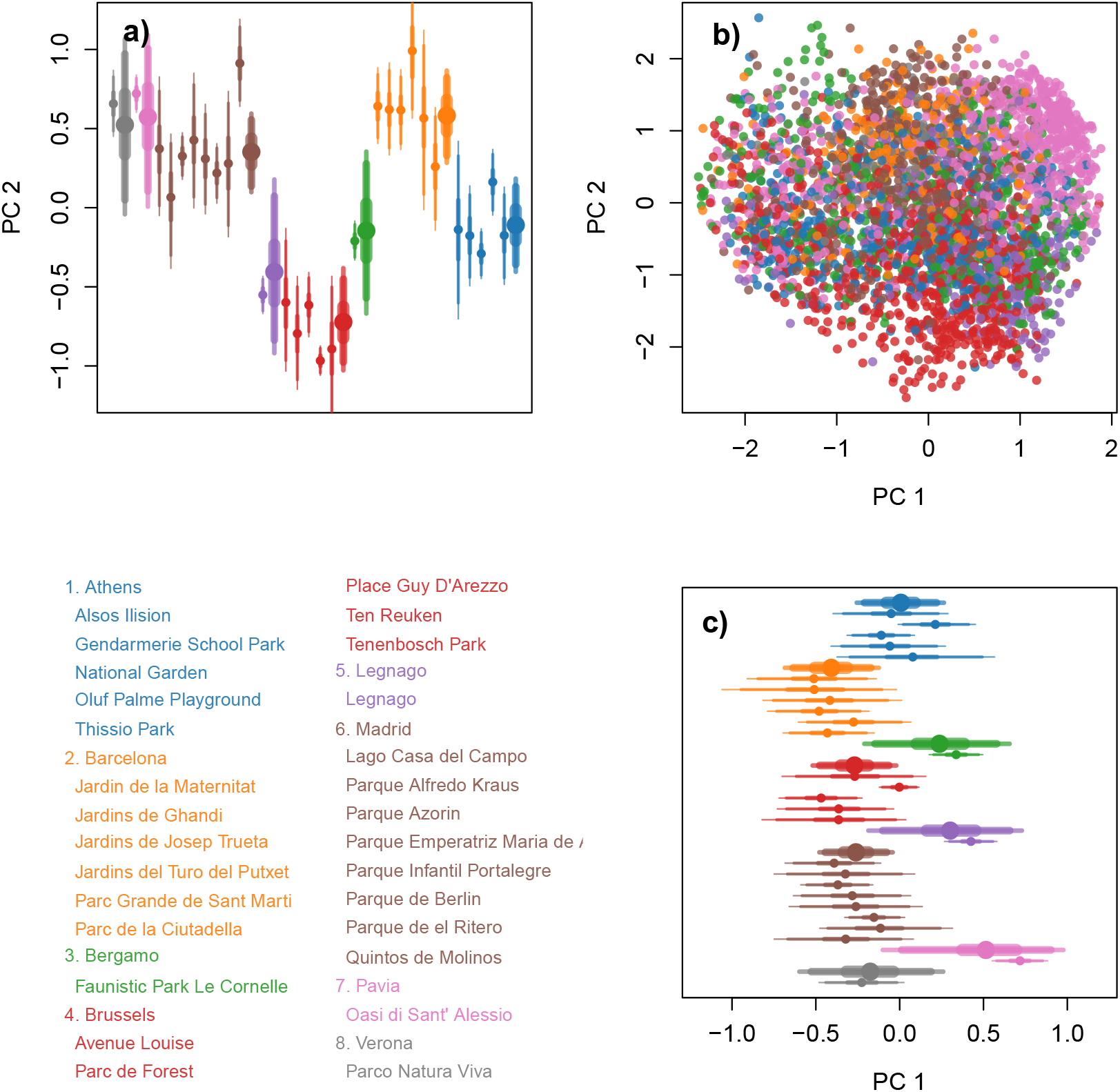
Result for PCO. Colour represents city (see legend). a) City (thick) and park (thin) averages (dots) and 50, 90 and 95% intervals for PC 2. b) Scatter-plot of all calls included in the model. c) City (thick) and park (thin) averages (dots) and 50, 90 and 95% intervals for PC 1.

In contrast, there was little evidence for widespread differences between parks within cities. Differences were only observed in a few cases. Lago Casa del Campo and Parque de el Ritero were clearly different from other parks in Madrid. Likewise, Gendarmerie School Park and the National Garden were different from other parks in Athens. It is important to mention that those observed park level differences could potentially be a result of incorrect pooling (i.e, assigning unique IDs to vocalisations from the same individual or assigning one ID to vocalisations from different individuals), as the standard deviation across parks was well within the values found in the sensitivity analysis (see Supplemental Materials, Figure S3). Park level means can appear very different under incorrect pooling, even when no signal exists in the simulated data (see Figure S1, Supplemental Materials). The city level signal we detected is much stronger than the simulated results due to incorrect pooling (see Supplemental Materials, Figure S2). This lends strong support for dialect differences between cities, while there is no support for this at the park level given the few differences observed.

In addition to assessing overall differences between parks and cities, we examined the proportion of contact call variants that were observed across the different cities (see Figure 5). We found that in most cities, the *typical* variant was predominant (see Figure 2a), and 4-5 other variants were usually present at intermediate to low frequencies. Multiple cities had a large proportion of contact calls that started with a low frequency component - *ladder start* (see Figure 2c). Pavia was characterized by the *four triangle* contact call with four triangular frequency modulations (see Figure 2b). Brussels stood out from the rest with the *mix alarm* contact call, containing multiple alarm-like notes (see Figure 2d).

**Figure 5.**
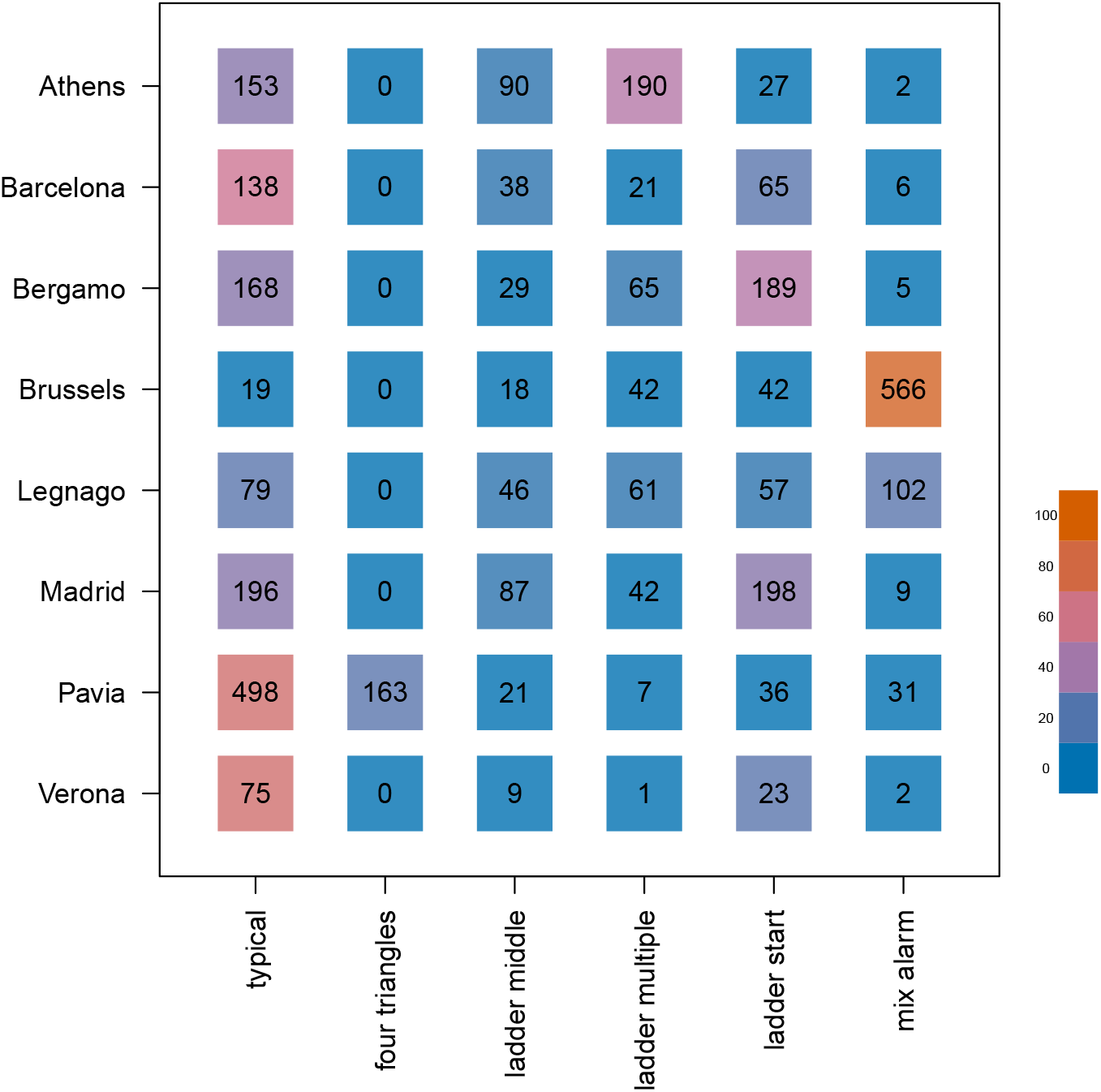
Distribution of variants across cities. Numbers are represented in black, colours represent the percentage of the given variant within the given city and range from 0% (blue) to 100% (orange) - see colour scale bar.

## Discussion

Our results provide strong evidence that monk parakeet contact calls differ between multiple cities across their European range. Vocal differences between the parks within cities were also detected, however, these differences were less consistent compared to the dialect pattern we observed at the city level and appeared to be only present in a few parks (see Figure 4). Overall, our results provide support for the *cultural drift hypothesis*, while finding no support for the *group membership hypothesis*. If vocal convergence was occurring at the group level, we would expect a stronger signal for dialects or clinal variation at the park level compared to city level, because movement between parks is likely very limited (Senar, Moyà, et al. 2021). Instead, our results demonstrate strong dialect differences at the city level. This result suggests that passive cultural processes are at play (Podos and Warren 2007; Sewall, Young, and Wright 2016; Bradbury and Balsby 2016). Finally, while we cannot directly test this hypothesis in our framework, the lack of consistent evidence for park level differences is a pattern in line with other monk parakeet research (Smith-Vidaurre, Araya-Salas, and Wright 2020) that found strong support for the *individual signature hypothesis*. We should note that this is not mutually exclusive with the *cultural drift hypothesis*. It could be that both are operating simultaneously at different spatial scales (Thomsen, Balsby, and Dabelsteen 2013), highlighting the importance of spatial scale in dialect studies.

Detecting the spatial scale at which geographic vocal variation emerges can be difficult, especially in a largely untagged population. For example, (Smith-Vidaurre, Araya-Salas, and Wright 2020) used partial Mantel tests and detected a signal at all scales of their analysis. However, they were not able to directly compare this to the individual signal, as sample sizes differed and Mantel tests do not provide a comparable statistic. A Bayesian multilevel model does provide such a statistic (*σ*_park_ and *s*_city_) and allows one to test the influence of incorrect pooling in a largely untagged population (see sensitivity analysis - Supplemental Materials). We can therefore say with a high degree of confidence that the city level signal outweighs the park level signal and is well above any spurious signal that might be due to incorrect pooling.

Previous studies in other parrot species have often argued that dialects arise at the group level because of selective pressures to conform to local variants (Wright and Dahlin 2018; Eberhard et al. 2022), including an active signalling of group membership. However, because we observed little evidence for dialects among parks, we do not think it likely that monk parakeets conform to local dialect types as a mechanism to identify group members. Instead, we find it more likely that the observed dialects among cities result from either random errors and conformity as described in the *cultural drift hypothesis*, or from an influence of the original founding populations (Ju et al. 2019). This supports other work in parrots that has also found dialects all be it at smaller geographical scales (Wright 1996; Buhrman-Deever, Rappaport, and Bradbury 2007; Martínez and Logue 2020; M. C. Baker 2003; Kleeman and Gilardi 2005).

Given the limited dispersal between European populations of monk parrots, another possibility is that there is vocal and genetic concordance, as is observed in crimson rosellas (Ribot et al. 2012) and palm cockatoos (Keighley et al. 2020). However, we find this unlikely in our study system. A previous study found that genetic differences between populations of monk parakeets in Europe are minimal, and that most areas were likely sourced from the same founding populations (Edelaar et al. 2015). Thus, genetic differences appear to be a less likely explanation for city level vocal differences than cultural processes, with the source groups determining the starting vocal dialect of each population. Even though previous work combined with our results suggest that monk parakeet contact calls are at least partially socially learned, the exact process is not fully resolved and the ontogeny of vocal learning needs more attention. It is well known that call structure of individuals is influenced by vertical transmission and the family environment (Berg, Delgado, Cortopassi, et al. 2012; Berg, Beissinger, and Bradbury 2013; Arellano et al. 2022). Prior research suggests that dispersing juveniles are the ones most likely to modify their calls after dispersal while adults do not (Wright and Dorin 2001). However, we did not observe clear dialects at the park level, to which juveniles could converge.

Interestingly, previous research on invasive monk parakeets suggests that dispersal between both parks and cities is very limited (Dawson Pell et al. 2021). Hence, we might expect cultural drift to also lead to dialects at the park level, yet we see the opposite pattern. Interestingly, we also found no support for clinal variation between parks (see further analysis in supplemental materials, where we tested the effect of distance on park-level vocal similarity). One possible explanation for why we do not observe dialects or geographic variation at the park level is provided by the *individual signature hypothesis*. Here, the lack of a clear park signature could be explained by divergence in order to stand out in acoustic space (Berg, Delgado, Okawa, et al. 2011). However, unlike the results from (Smith-Vidaurre, Araya-Salas, and Wright 2020), which suggest that selection for individually distinctive calls outweighs any selection for call convergence at the group level, we found very clear evidence for dialects between cities. A possible explanation for this discrepancy is that the study undertaken by Smith-Vidaurre, Araya-Salas, and Wright (2020) was undertaken in the native distribution of monk parakeets, while our results were obtained in a large invasive range where populations are fragmented and dispersal between populations (i.e., cities) is very unlikely (Dawson Pell et al. 2021; Bucher, Martin, et al. 1990). In contrast, although dispersal patterns have not been fully described in the native range, the habitat is more continuous, with increases in Eucalyptus trees allowing for long distance dispersal across the entire range (Bucher and Aramburú 2014; Da Silva et al. 2010). Furthermore, monk parakeets are considered an agricultural pest and are heavily persecuted in their native range (Castro, Sáez, and Molina-Morales 2021). The effect of persecution is often increased dispersal and between-group movement (Payo-Payo et al. 2018) leading to increased intermixing between sub-populations that could potentially obscure any dialect patterns. Such differences in dispersal might partially explain why dialects were also detected in populations of invasive monk parakeets in the United States (Buhrman-Deever, Rappaport, and Bradbury 2007).

While we did not find evidence for strong convergence towards a group level signature in contact calls, it could be the case that group signatures exist in other call types, or within very specific variants of contact calls. In accordance to our call type analysis, (see Figure 5), most variants were present in all cities, but some showed higher proportions than others. While we cannot be certain that these variants drive the dialect differences between cities, or lack of in parks, they raise an important point. Explicit experiments that strive to determine the function of these can help us understand where and when to expect the stronger variation between them. Further complicating this, is that as vocal learners, it is possible that certain populations learn to use different variants in different contexts. The ontogeny of these variants, as well as the contextual mechanisms will help further the study of dialect mechanisms in not only Monk Parakeets, but all Psitticine species. In (Wright and Dorin 2001), it was found that juvenile birds more readily modified their contact calls after translocation than adult birds. Given that our populations started from invasive released birds, it could be a critical piece of information to know what the population dynamics were at the beginning of invasion, and the dynamics of subsequent invasion.

An alternative explanation for the lack of strong park signals could be that group signatures exist at a smaller scale. Monk parakeets nest in complex nest structures and previous work has shown that birds from the same nest tree are more closely related than expected by chance and tend to forage together (Dawson Pell et al. 2021). This might suggest that either passive or active processes could instead result in a nest level, rather than park level, signature. Future studies should focus on a single population and estimate the strength of the individual and group level signatures across multiple scales. This should preferentially be done in an individually-marked population, such that the temporal stability of vocalisations can also be estimated. Lastly, we recommend that playback studies be conducted on monk parakeet across populations at both the park and city level to indeed experimentally test whether birds can detect subtle variations in group signatures, not picked up by our analyses. For example, tests could examine whether birds recognize calls from their own versus distant colonies, as well as other cities. Furthermore, playback tests could be used to test different substructures of the park (i.e., family unit, specific tree) to see if the park scale is an appropriate scale to measure vocal variation. This type of research is needed before dismissing the *group membership hypothesis* as a possible mechanism.

Geographic vocal variation is one of the primary forms of evidence for vocal learning (Lemon 1975; Marler and Tamura 1962). However, our understanding of the processes that lead to this variation at different scales and levels of population structure is lacking. A thorough understanding of these processes is critical to elucidating the underlying mechanisms that drive vocal learning and dialect formation. Monk parakeets and other parrot species are particularly useful model species to study social dynamics and vocal learning because of their flexible learning and complex social system. By continuing to apply novel techniques to the study of vocal patterns at different scales, we can uncover more detailed mechanisms of how communication systems evolve in natural populations. Our study demonstrates the existence of distinct dialects in European populations of monk parakeets, lending support to the *cultural drift hypothesis* while simultaneously showing patterns inconsistent with the *group membership hypothesis*. In addition to cultural drift, we also found evidence consistent with the *individual signature hypothesis* at the park level. While further experimental study is needed to confirm or refute these hypotheses, our extensive dataset, broad geographic scope and two-level comparison provide critical and robust information that enhances our understanding of the important role vocal learning plays in generating dialect differences among populations of Psittacine species.

## Supporting information

sensitivity analysis

## Ethics

All data was collected without disturbing the animals and, therefore, this study did not require any permits.

## Data, code and materials

All small data files and code are publicly available on https://github.com/simeonqs/Multi-level_Bayesian_analysis_of_monk_parakeet_contact_calls_shows_dialects_between_European_cities.

Large data files are available to reviewers upon request. All data and code will be stored permanently on Edmond upon acceptance.

## Acknowledgements

We would like to acknowledge Dr. Silke Atmaca and Dr. Bret A. Beheim for their research coordination assistance. We are very grateful to the help from Nina Schwarz, Philine Adolphi, Vivien Kleinow, and Gustavo Alarcón-Nieto for their assistance processing and organizing data. Thank you to Natagora, Research Department (Alain Paquet) in Brussels, Belgium for his guidance in finding populations of Monk Parakeets to record. Thank you to Roberta Castiglioni at Parco Faunistico Le Cornelle, Giulio Salamon at Oasi di Sant’Alessio, Caterina Spiezio at Parco Natura Viva, and Dr. Juan Carlos Senar at Cituadella Park for all their assistance and permission to record parakeets at these locations. We would like to thank Dr. Robert F. Lachlan for his guidance in using Luscinia. Finally, we would like to thank Prof. Jack W. Bradbury, Dr. Susannah Buhrman-Deever and Dr. Grace Smith-Vidaurre for valuable advice during the early phases of this project.

## Author contributions

Conceptualization: SQS, SAT, LMA, MBM; Data curation: SQS, SAT; Formal analysis: SQS, SAT; Funding acquisition: LMA, MBM; Investigation SQS, SAT; Methodology: SQS, SAT; Project administration: SQS, SAT; Resources: LMA, MBM; Software: SQS, SAT; Supervision: LMA, MBM; Validation: SQS, SAT; Visualization SQS; Writing – original draft: SQS, SAT; Writing – review & editing: SQS, SAT, LMA, MBM.

