## Supplementary material for "Multi-level Bayesian analysis of monk parakeet contact calls shows dialects between European cities": sensitivity analysis

#### 1 Justification

The aim of this sensitivity analysis was to test how our labelling of individuals affected the parameter estimates. We could only assign a subset of the vocalisations to specific birds and could not be certain that we did not rerecord these individuals later on. The remaining vocalisations were grouped under a unique ID for each five minute chunk in the recording. We therefore wanted to test the effect of three potential errors: 1) focal individuals could be the same across recordings, 2) calls coming from different five minute chunks could belong to the same individual, 3) calls within a five minute chunk could come from different individuals. The third error would not inflate the park and/or city level effect, since it would only increase heterogeneity at the individual level. The first and second error are potentially much more problematic.

#### 2 Methods

To test the combined effect of all three errors we ran multiple simulations where we varied the strength of the three errors. We used the same data structure as in the real dataset (same number of cities, parks per city, recordings per park and calls per recording) and kept verbally annotated IDs wherever available. We compared these simulations to a simulation where it was assumed that the IDs were known. In all simulations the standard deviation within and between individuals was 1 and there was no effect of either park or city. To test the effect of the three types of errors we included the following parameters:

1) lambda-rerecord:

This parameter ranged from 1 to 2 and was used to generate a poison distribution from which the number of recordings per individual was drawn. If this number exceeded 1, another ID from the same park was replaced with the ID of the focal individuals. This was done iteratively, until all individuals had either been replaced or been considered as focal individual. The new set of IDs was then used to generate individual level means. The original set of IDs was used during analysis of the simulated data.

2) p-next-chunk:

This parameter ranged from 0 to 0.7 and was used as probability for recording the same individual in the next five minute chunk. For each five minute chunk (which had been assigned a unique ID) we sampled whether or not it should be assigned the same ID as the previous chunk with the p-next-chunk determining the probability of drawing a TRUE. This was done iteratively for all chunks and the new IDs were used in a similar manner as for lambda-rerecord.

3) lambda-ind-per-rec:

This parameter was set to 1.5 and was used to generate a poison distribution from which the number of individuals per five minute chunk was drawn. Since this parameter should not affect the estimation of the park and city level effect, we did not run a full sensitivity analysis, but instead fixed it to the value that we thought most appropriately. For the vocalisations where the five minute chunks were used to generate IDs, we drew a value from this distribution. If the value exceeded 1, we reassigned the vocalisations to n new unique IDs. These new IDs were then used in a similar manner as for lambda-rerecord.

### 3 Results

Mislabelling had a strong effect on the park level estimates. Even low error rates inflated the posterior distribution of sigma_park (see Figure 1).

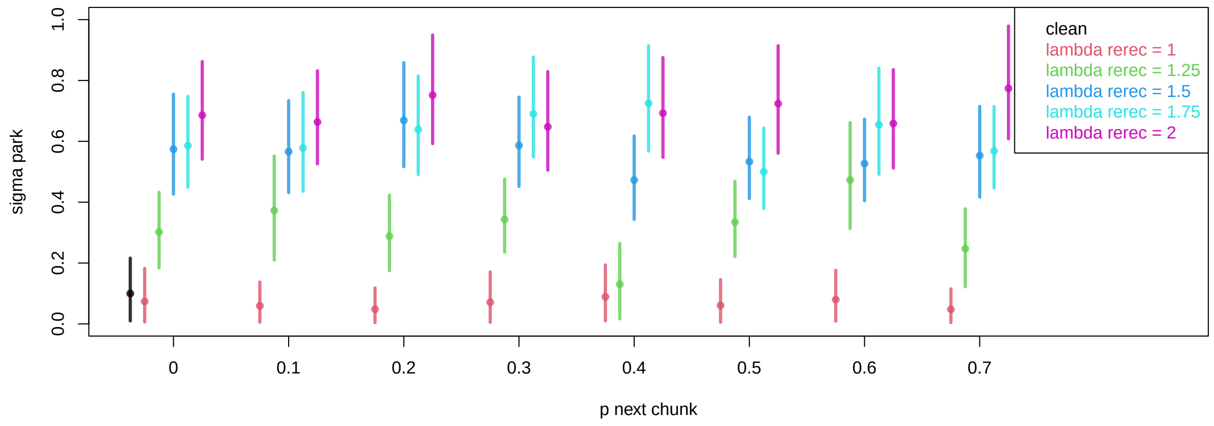


Figure 1: *Park level effect. Dots represent means of the posterior distributions. Lines represent 89% compatibility intervals.*

Mislabelling had very little effect on the city level estimates. Means were inflated, but so were the 89% compatibility intervals (see Figure 2), meaning that spurious effects would be detectable.

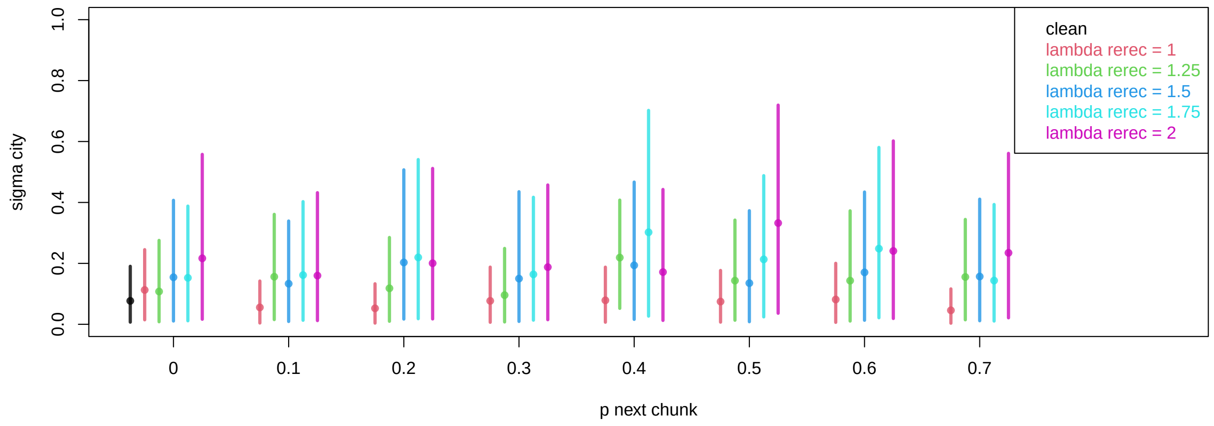


Figure 2: *City level effect. Dots represent means of the posterior distributions. Lines represent 89% compatibility intervals.*

To visualise the result of mislabelling on the city and park level means, Figure 3 displays how the data and model results for lambda-rerecord = 1.5 and p-next-chunk = 0.4. Parks are clearly different. Cities are also somewhat different, but the effect is much weaker than in the real data.

The sigma parameters for the real data were within what could be a spurious result for the park level effect, but were much farther from 0 for the city level effect (see Figure 4).

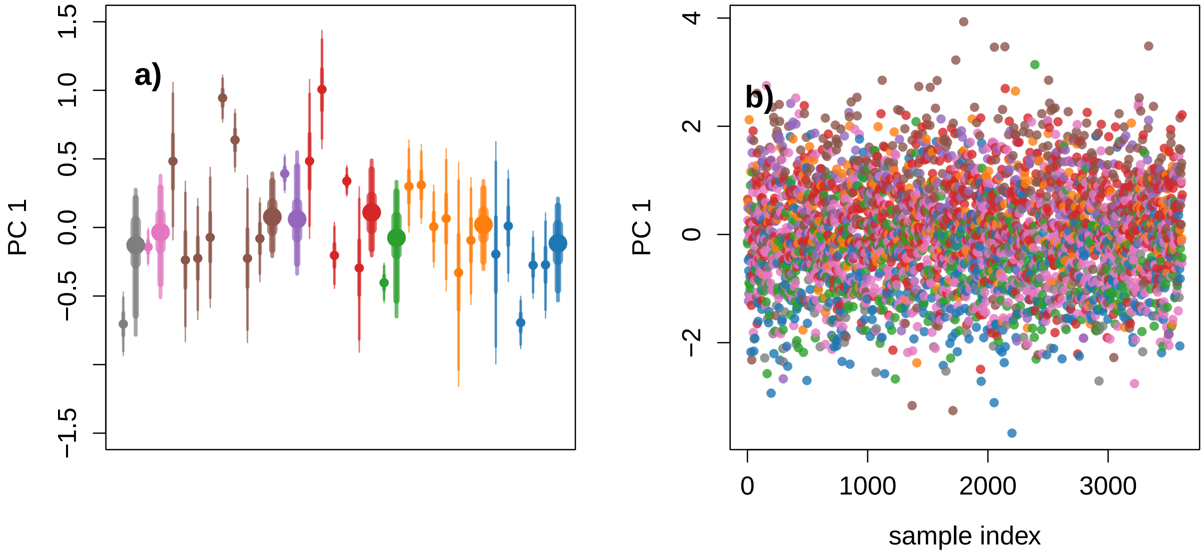


Figure 3: *Effect of mislabeling on city and park level means.* **a)** *City (thick) and park (thin) averages (dots) and 50, 90 and 95% intervals (lines) for simulated PC 1.* **b)** *Scatter-plot of all simulated calls included in the model.*

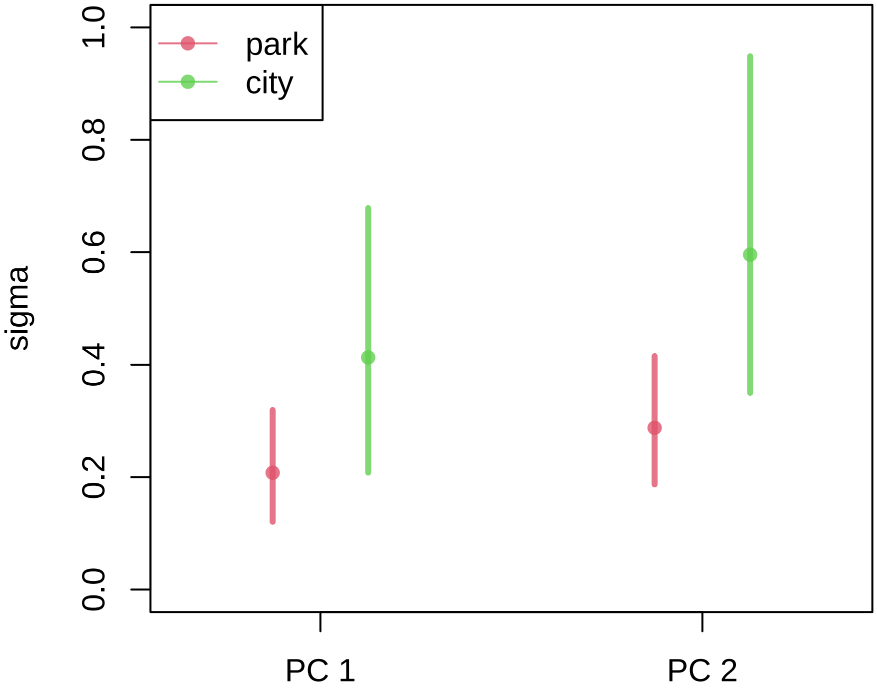


Figure 4: *Sigma estimates for the real data. Averages (dots) and 89% intervals (lines).*
