## supplemental materials for "Multi-level Bayesian analysis of monk parakeet contact calls shows dialects between European cities"

### 1 Methods

Contact call similarity was measured using three different methods:

1) dynamic time warping on fundamental frequency traces from Luscinia (DTW): see main text
2) cross correlation on fundamental frequency traces from Luscinia (CC): a custom written script (see ANALYSIS/CODE/function/simple.cc.R) was used to slide two traces over each other and measure the pointwise absolute difference. The minimal value (maximal overlap) was standardised by the length of the longest trace and used as similarity measure.
3) spectrographic cross correlation (SPCC): for each call we applied a high-pass filter using *ffilter* from the package *seewave* (Sueur, Aubin, and Simonis 2008) with settings: from = 500 Hz. A custom written script was then used to create spectrograms (see ANALYSIS/CODE/function/cutte.spectro.R). We used *specgram* from the package *signal* (signal developers 2014) with window = 512 and overlap = 450 to create the basic spectrogram. Only the 1-6 kHz range was included in the final spectrogram. All pixels were standardised. All pixels with a value lower than 1.3 were then set to 0 and pixels with a value higher than 1.8 were set to 1.8. This was done to remove as much noise as possible and reduce the effect of particularly loud sections of the call. As a final step each pixel was divided by the summed value of all pixels, making sure that each spectrogram summed to one. This was done to make long and short call comparable.

Dimensionality of the resulting distance matrices was reduced to two using three methods:

1) principle coordinate analysis (PCO): see main text
2) principle component analysis (PCA): using *princomp* from the basic *stats* package in R
3) uniform manifold approximation and projection (UMAP): using *umap* from the package *umap* (Konopka 2020) with the settings: input = 'dist', n_neighbors = 10, spread = 1, min_dist = 0.1

For all possible combinations of similarity measures with dimension reduction we ran the same Bayesian multilevel model on the two remaining dimensions (see main text).

#### 1.1 Distance

To test the effect of distance between parks we ran a Bayesian model similar to the model in the main text, but within each city. We used a multi-normal distribution where the variance-covariance was computed from the distance matrix using the L2-norm.

### 2 Results

Figures 1-9 show the composite figures for all methods.

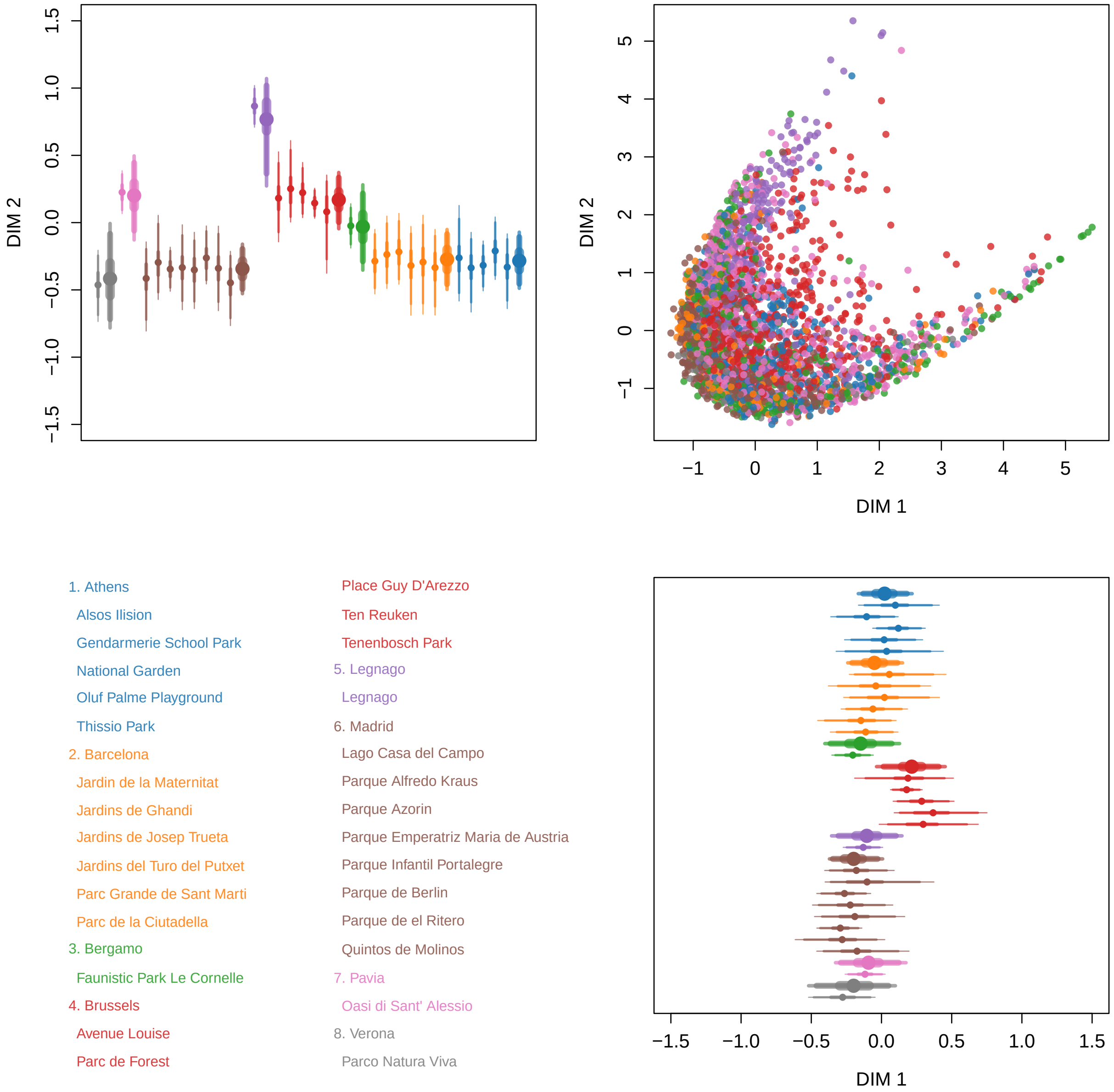


Figure 1: Result for CC - PCA. Colour represent city (see legend). Clock-wise: City (thick) and park (thin) averages (dots) and 50, 90 and 95% intervals for DIM 1; Scatter-plot of all calls included in the model; City (thick) and park (thin) averages (dots) and 50, 90 and 95% intervals for DIM 2.

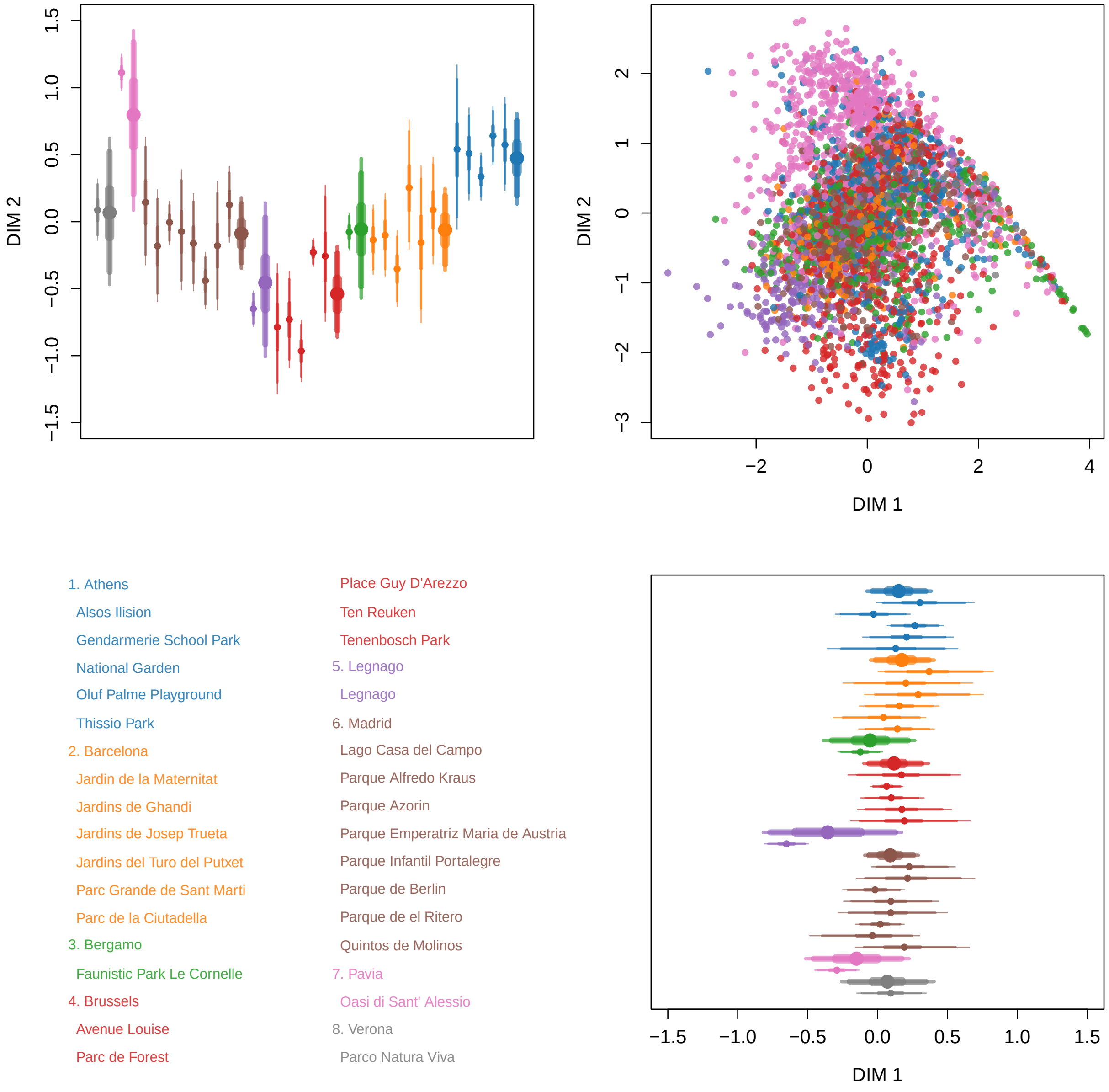


Figure 2: Result for CC - PCO. Colour represent city (see legend). Clock-wise: City (thick) and park (thin) averages (dots) and 50, 90 and 95% intervals for DIM 1; Scatter-plot of all calls included in the model; City (thick) and park (thin) averages (dots) and 50, 90 and 95% intervals for DIM 2.

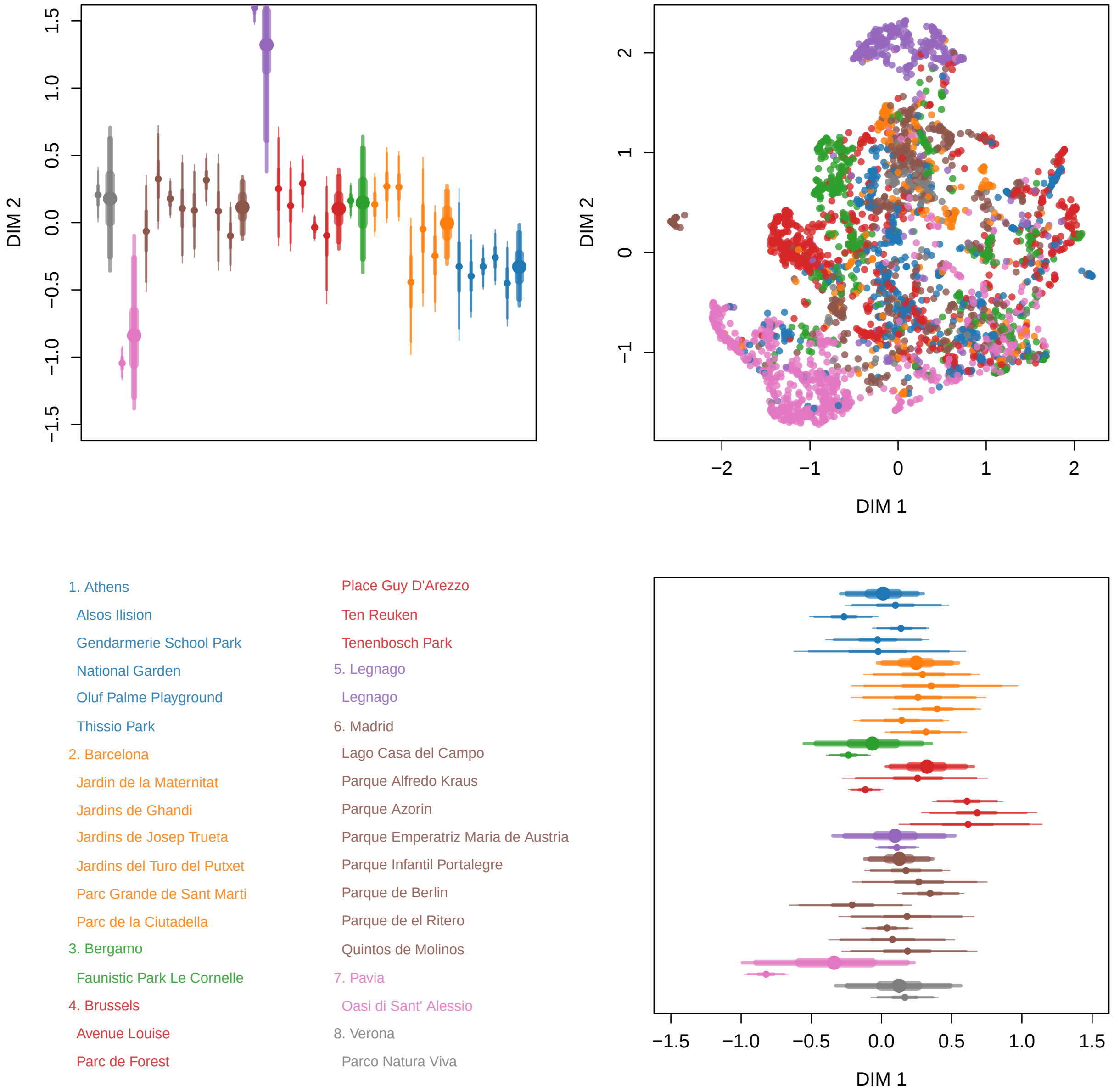


Figure 3: Result for CC - UMAP. Colour represent city (see legend). Clock-wise: City (thick) and park (thin) averages (dots) and 50, 90 and 95% intervals for DIM 1; Scatter-plot of all calls included in the model; City (thick) and park (thin) averages (dots) and 50, 90 and 95% intervals for DIM 2.

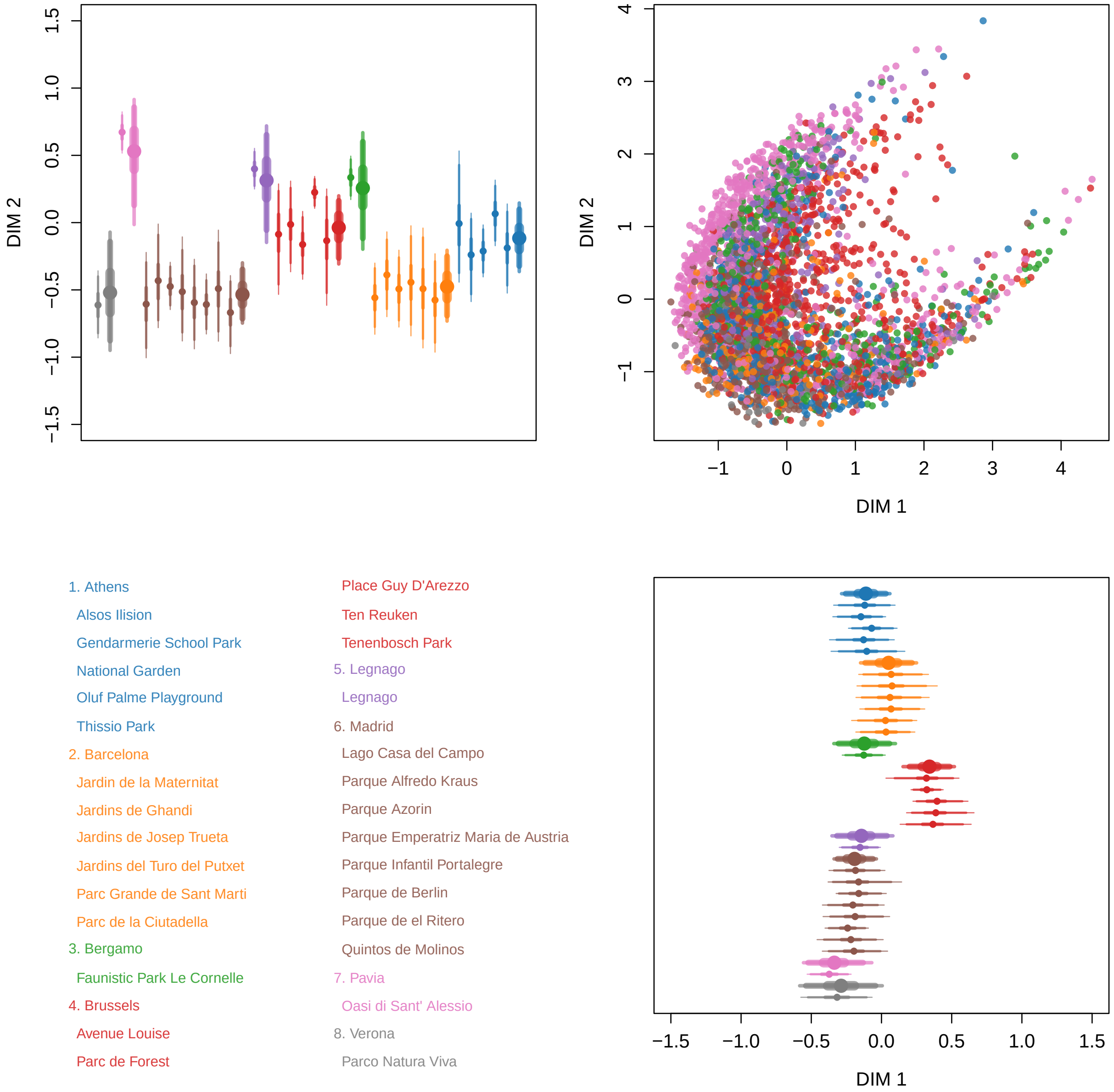


Figure 4: Result for DTW - PCA. Colour represent city (see legend). Clock-wise: City (thick) and park (thin) averages (dots) and 50, 90 and 95% intervals for DIM 1; Scatter-plot of all calls included in the model; City (thick) and park (thin) averages (dots) and 50, 90 and 95% intervals for DIM 2.

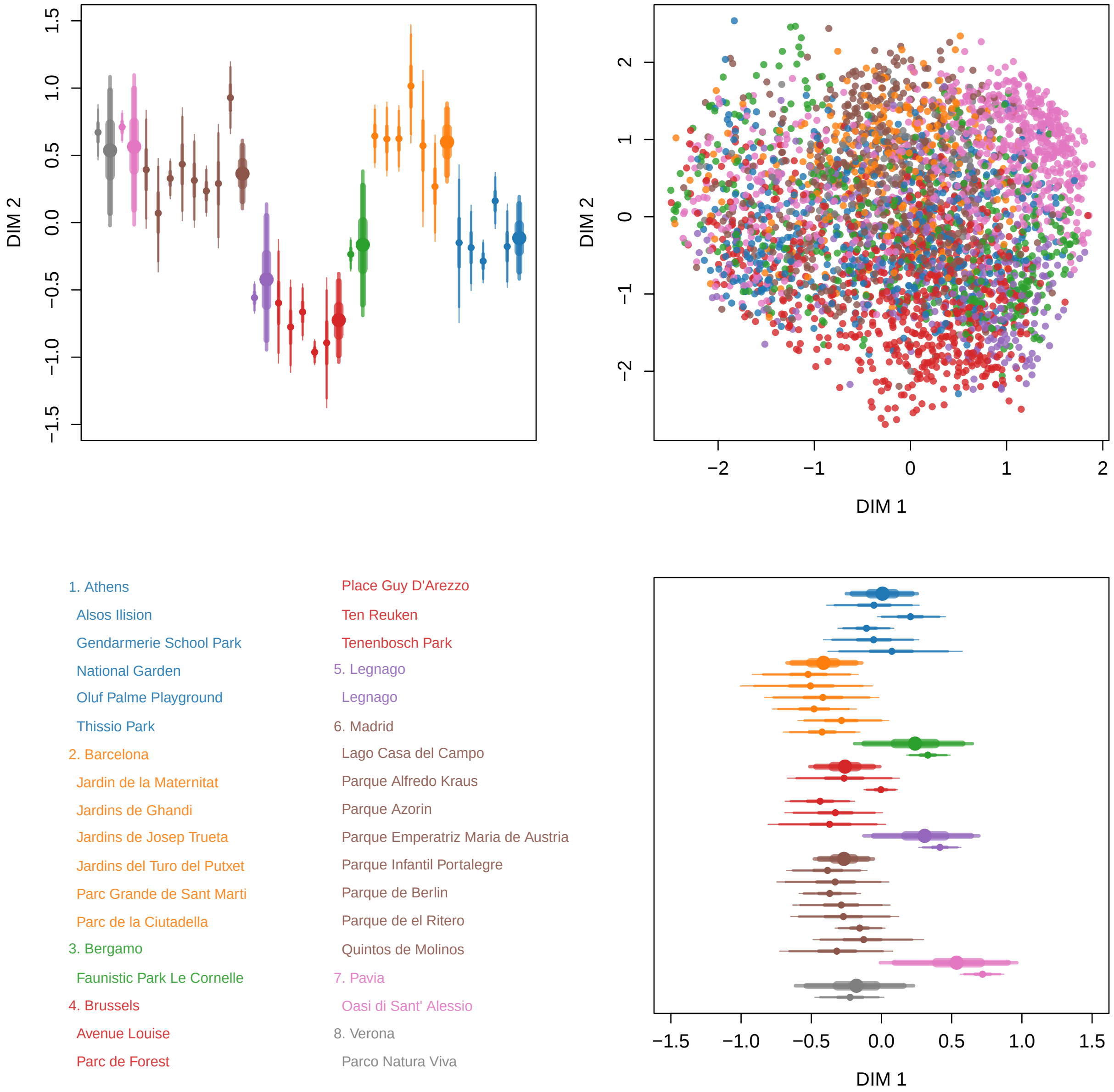


Figure 5: Result for DTW - PCO. Colour represent city (see legend). Clock-wise: City (thick) and park (thin) averages (dots) and 50, 90 and 95% intervals for DIM 1; Scatter-plot of all calls included in the model; City (thick) and park (thin) averages (dots) and 50, 90 and 95% intervals for DIM 2.

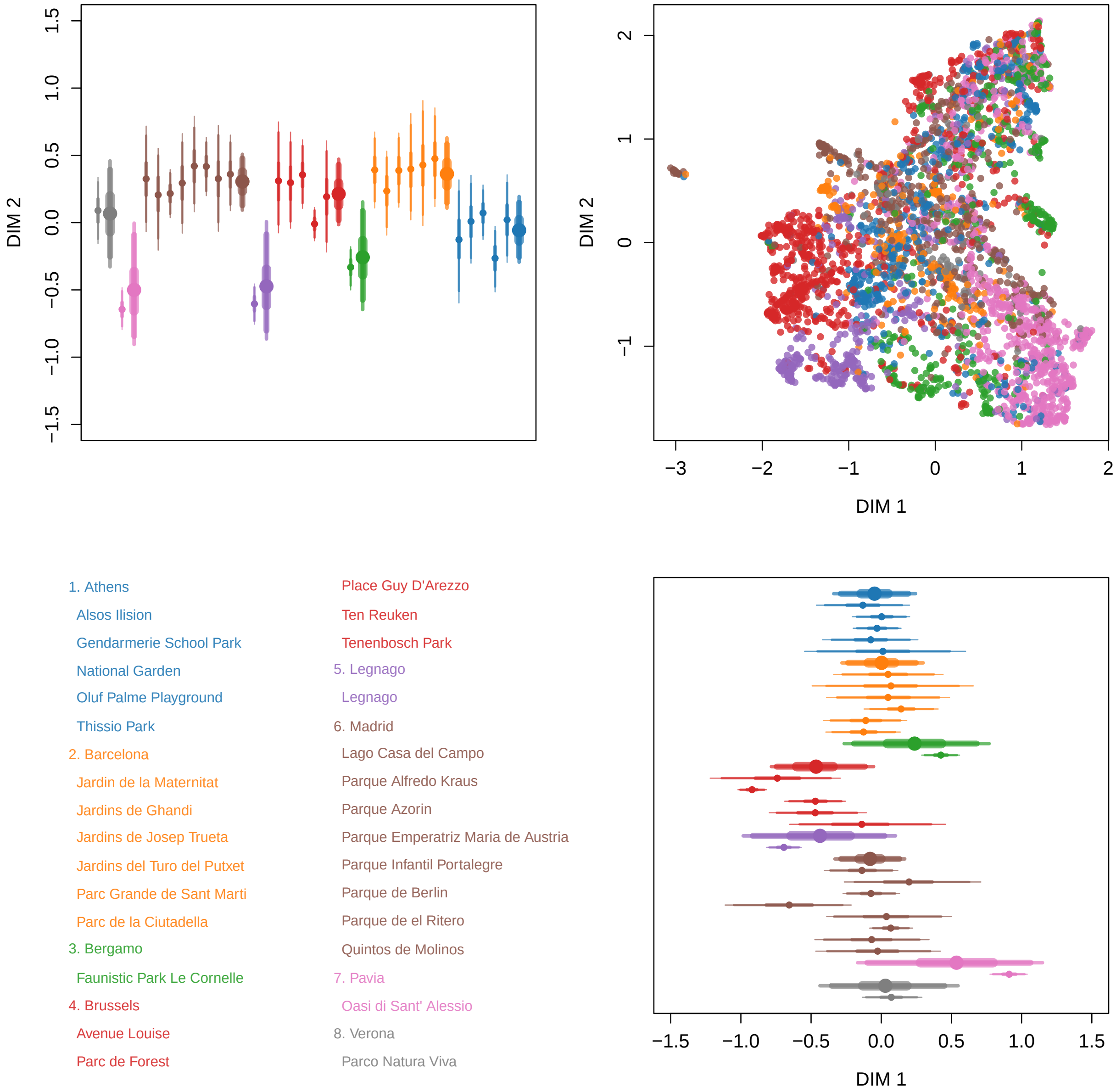


Figure 6: Result for DTW - UMAP. Colour represent city (see legend). Clock-wise: City (thick) and park (thin) averages (dots) and 50, 90 and 95% intervals for DIM 1; Scatter-plot of all calls included in the model; City (thick) and park (thin) averages (dots) and 50, 90 and 95% intervals for DIM 2.

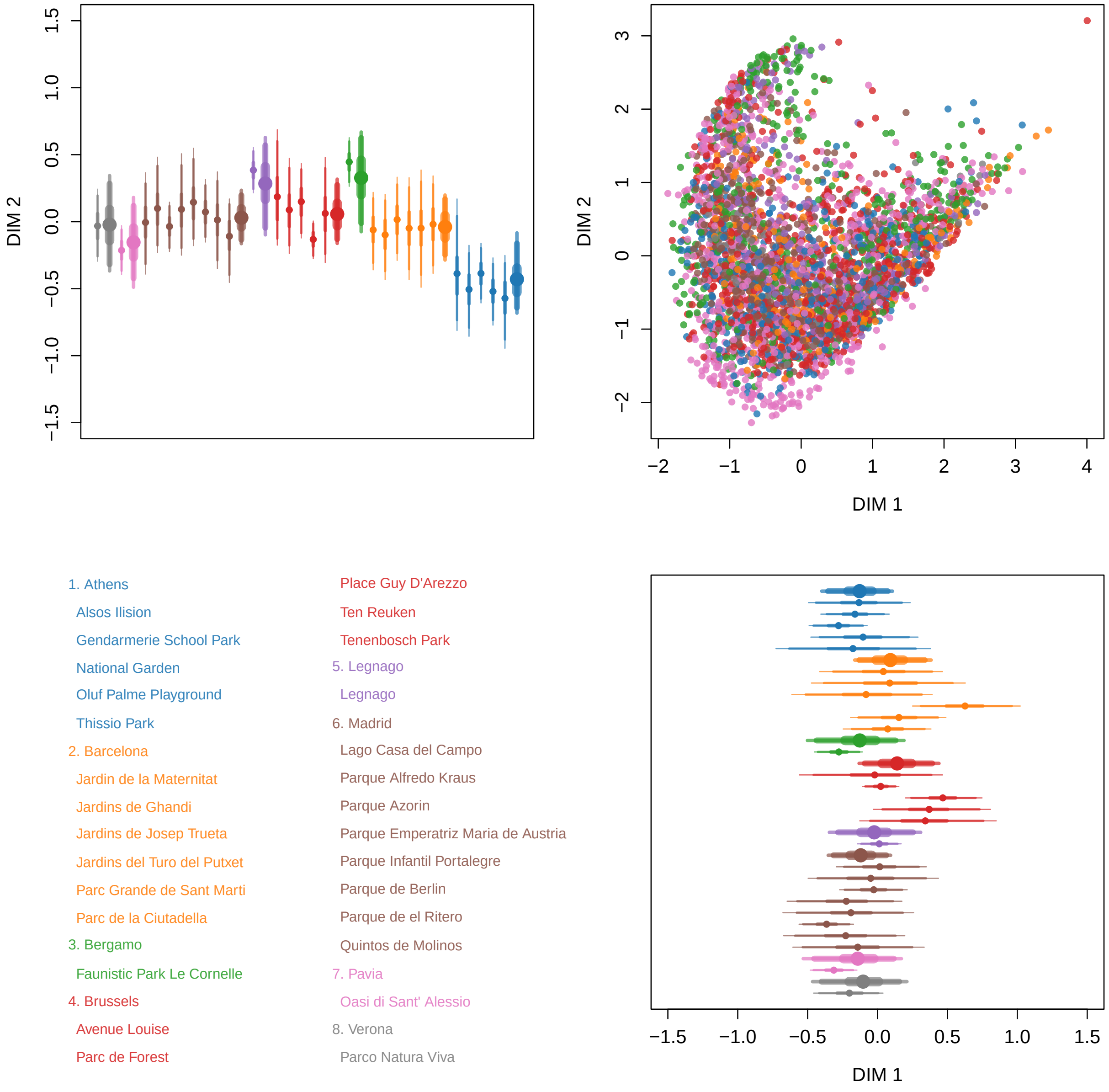


Figure 7: Result for SPCC - PCA. Colour represent city (see legend). Clock-wise: City (thick) and park (thin) averages (dots) and 50, 90 and 95% intervals for DIM 1; Scatter-plot of all calls included in the model; City (thick) and park (thin) averages (dots) and 50, 90 and 95% intervals for DIM 2.

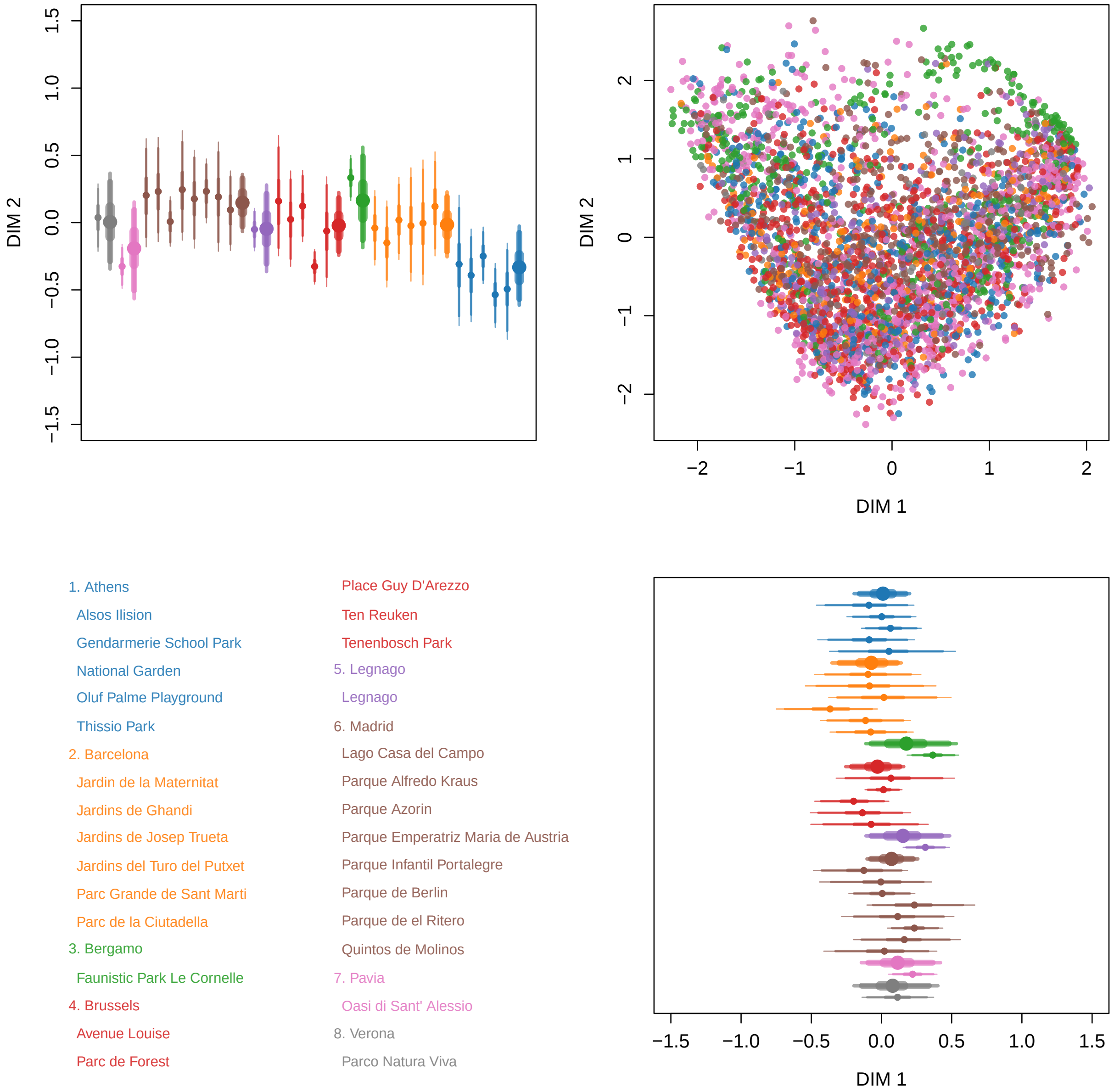


Figure 8: Result for SPCC - PCO. Colour represent city (see legend). Clock-wise: City (thick) and park (thin) averages (dots) and 50, 90 and 95% intervals for DIM 1; Scatter-plot of all calls included in the model; City (thick) and park (thin) averages (dots) and 50, 90 and 95% intervals for DIM 2.

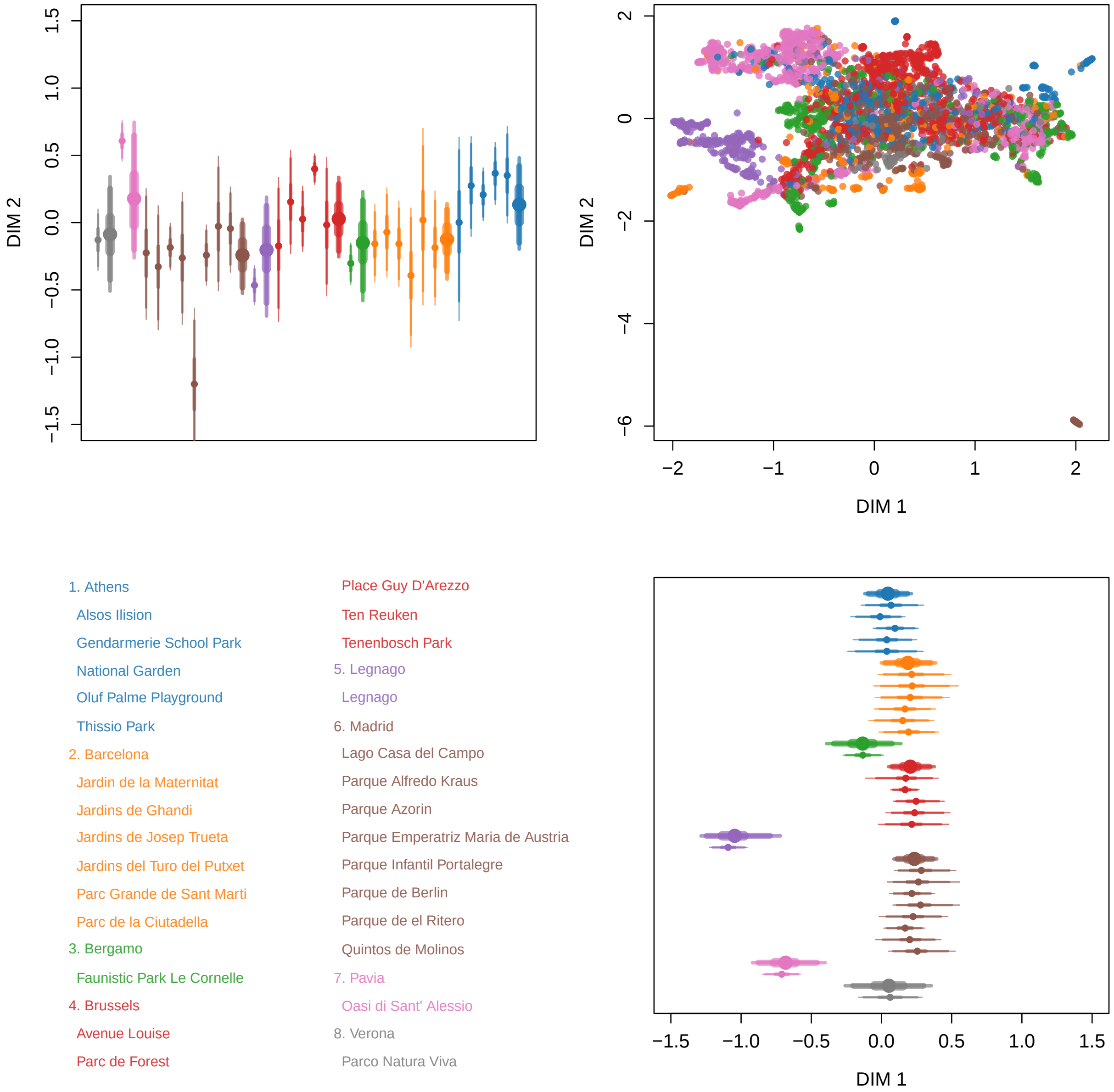


Figure 9: Result for SPCC - UMAP. Colour represent city (see legend). Clock-wise: City (thick) and park (thin) averages (dots) and 50, 90 and 95% intervals for DIM 1; Scatter-plot of all calls included in the model; City (thick) and park (thin) averages (dots) and 50, 90 and 95% intervals for DIM 2.

Figures 10-18 show the sigma values for all methods.

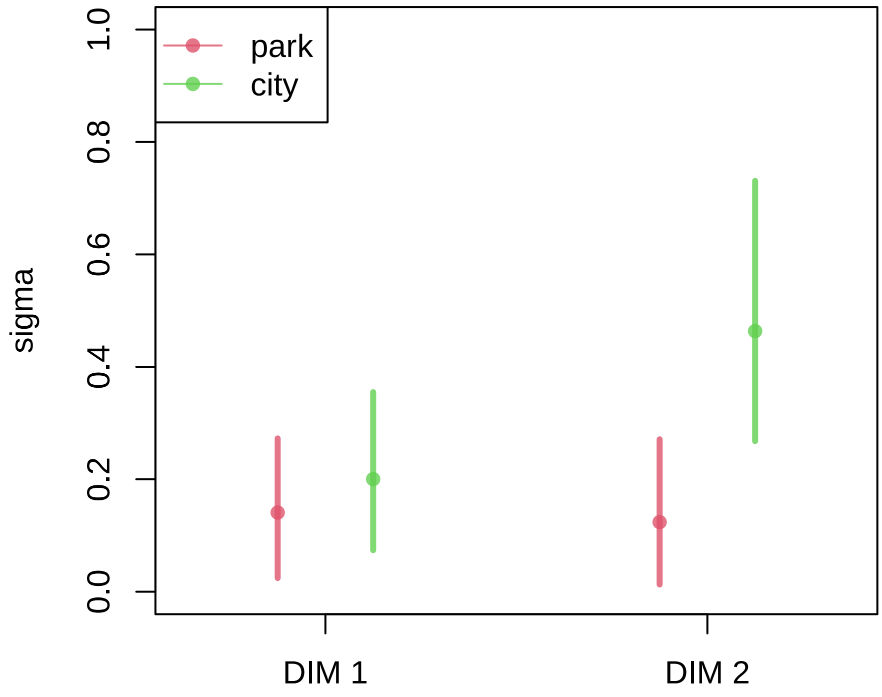


Figure 10: Result for CC - PCA. Sigma values for city and park.

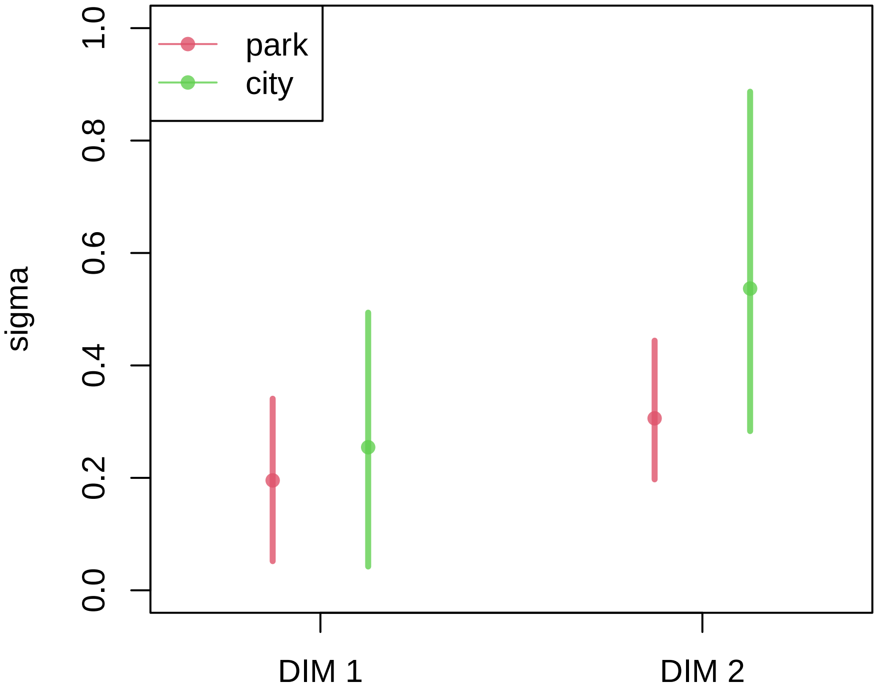


Figure 11: Result for CC - PCO. Sigma values for city and park.

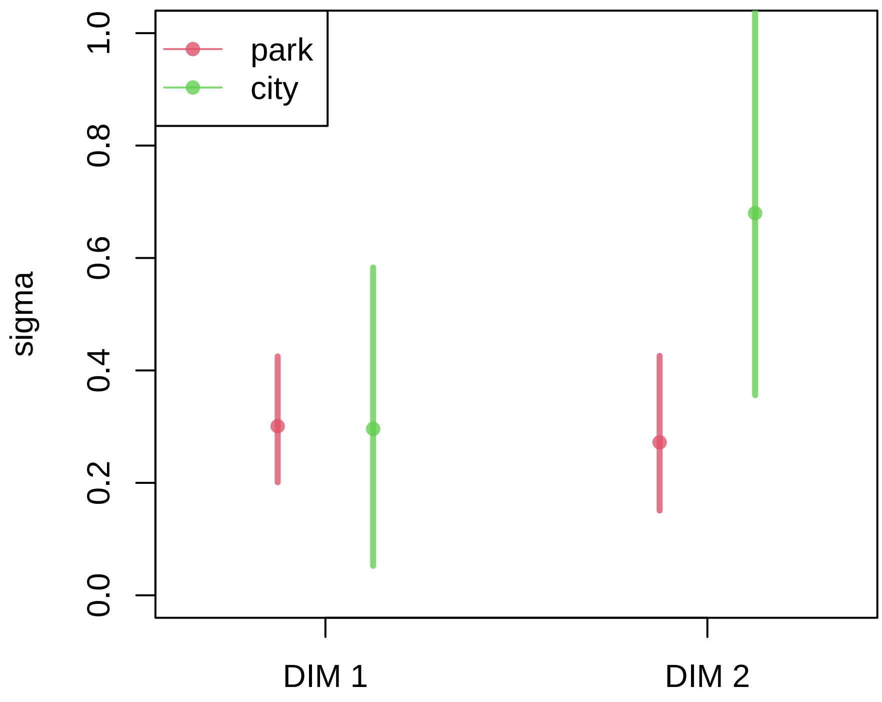


Figure 12: Result for CC - UMAP. Sigma values for city and park.

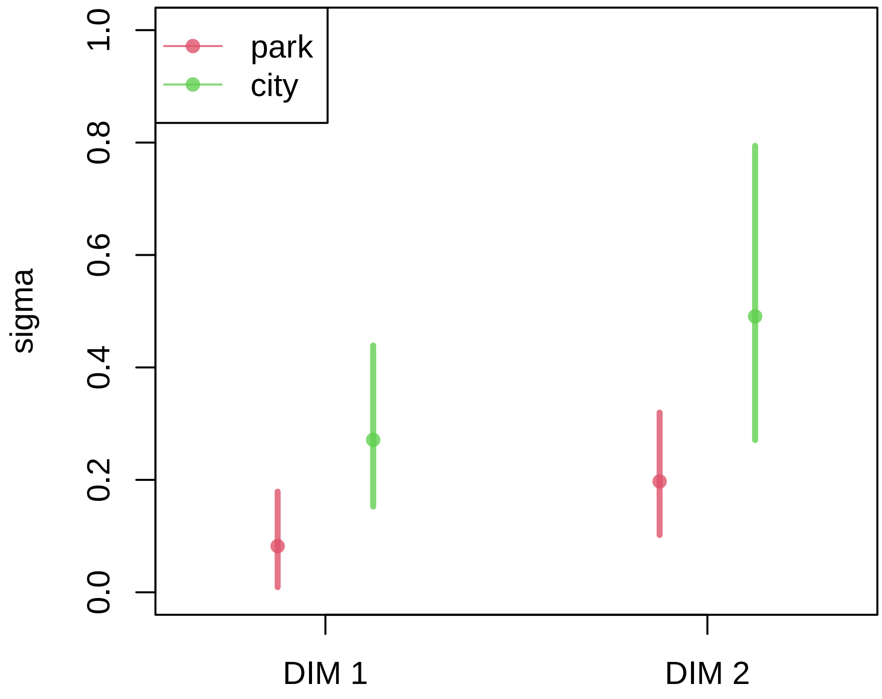


Figure 13: Result for DTW - PCA. Sigma values for city and park.

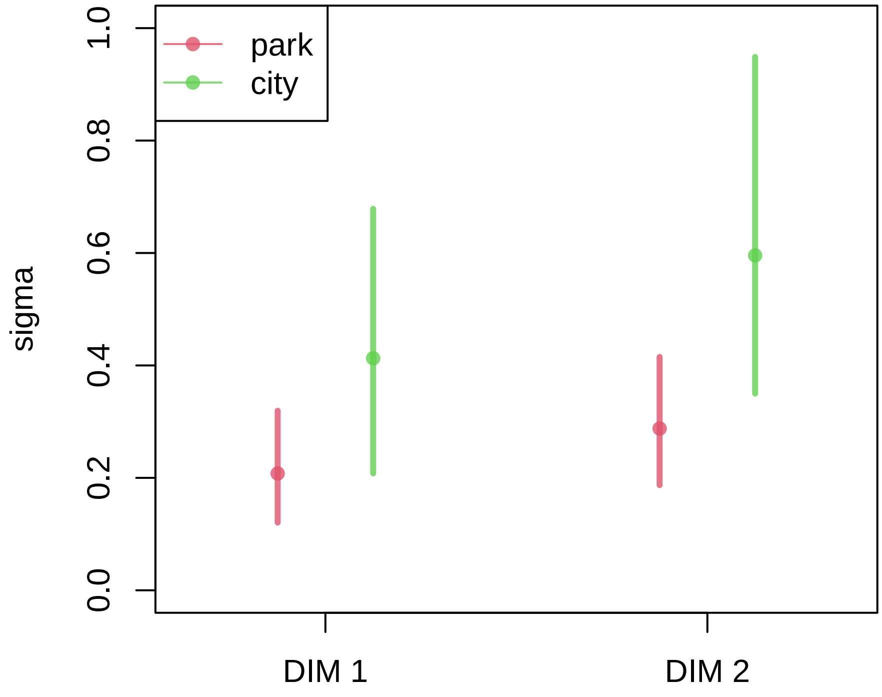


Figure 14: Result for DTW - PCO. Sigma values for city and park.

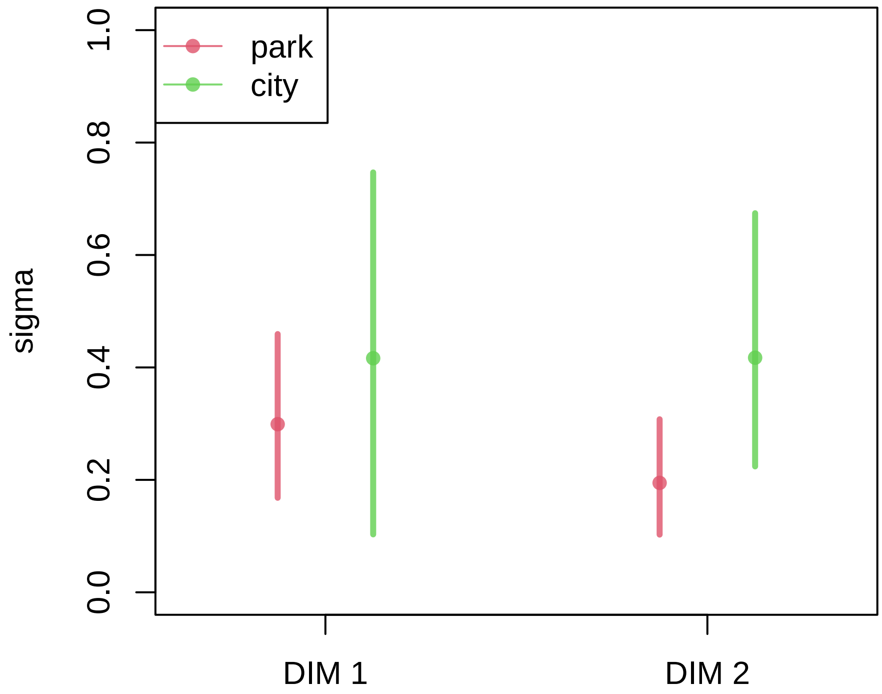


Figure 15: Result for DTW - UMAP. Sigma values for city and park.

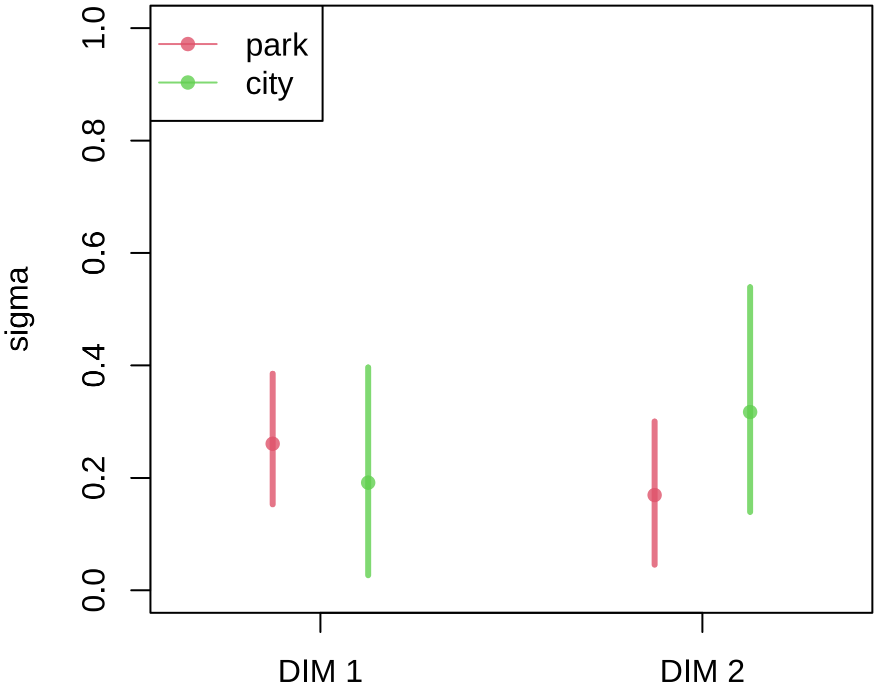


Figure 16: Result for SPCC - PCA. Sigma values for city and park.

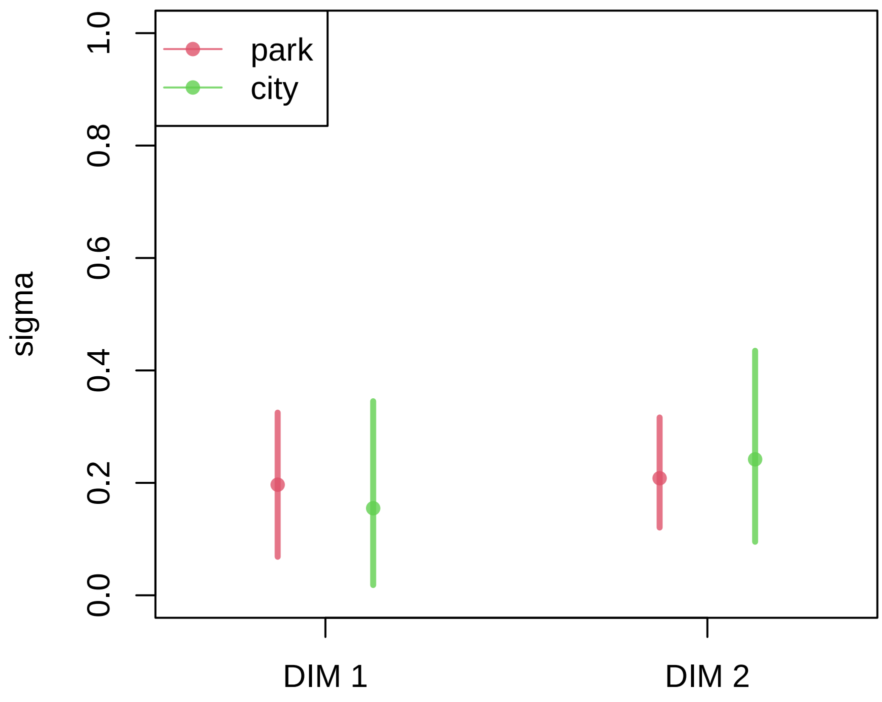


Figure 17: Result for SPCC - PCO. Sigma values for city and park.

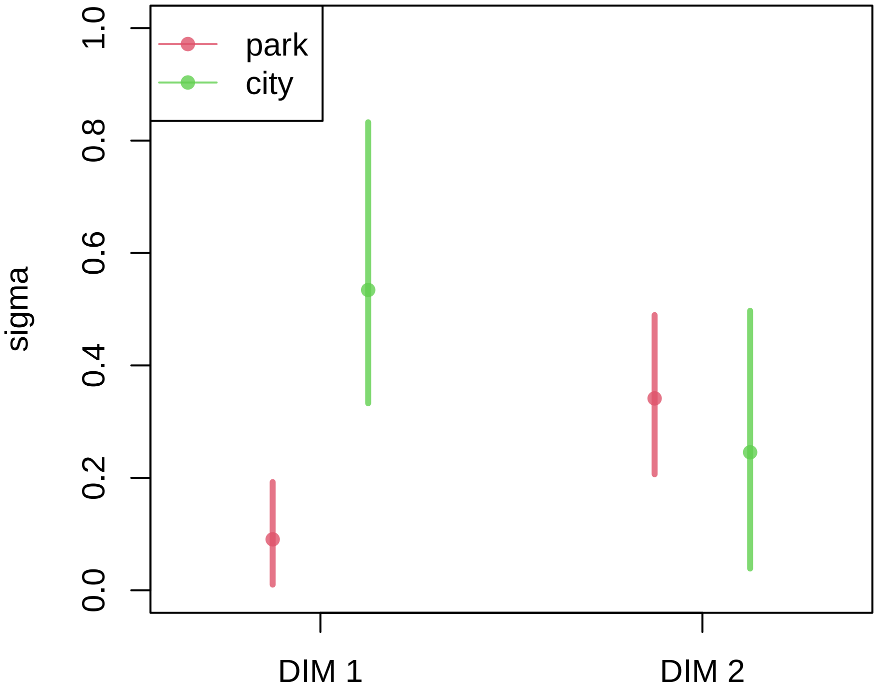


Figure 18: Result for SPCC - UMAP. Sigma values for city and park.

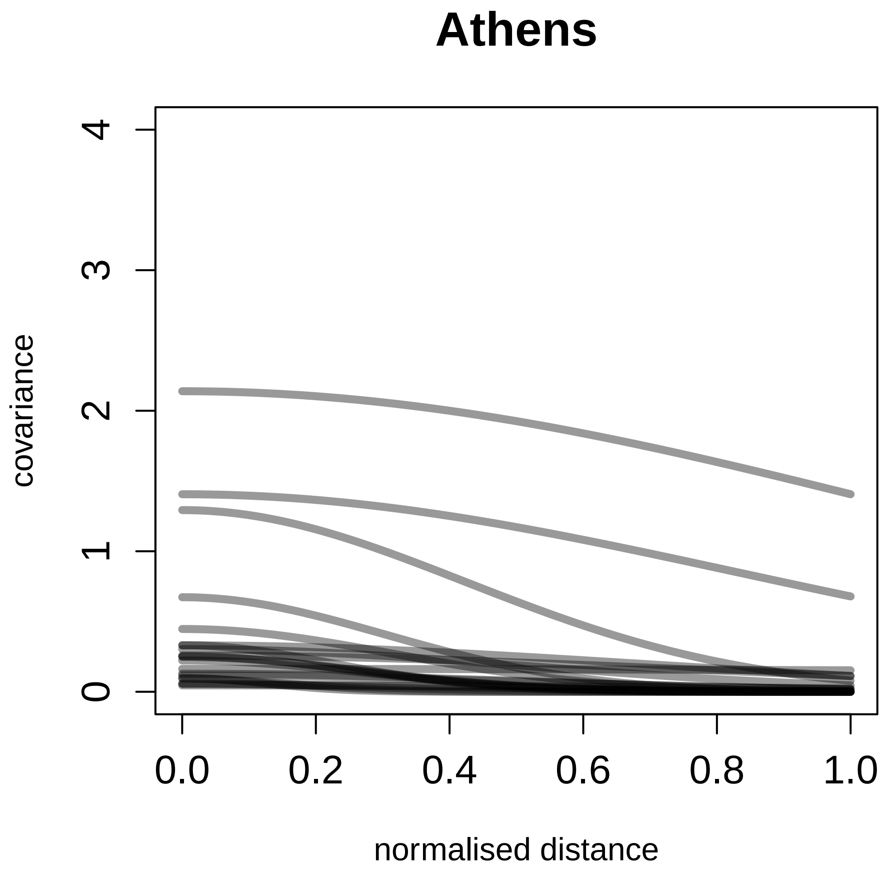


Figure 19: Covariance between parks in Athens as function of normalised distance.

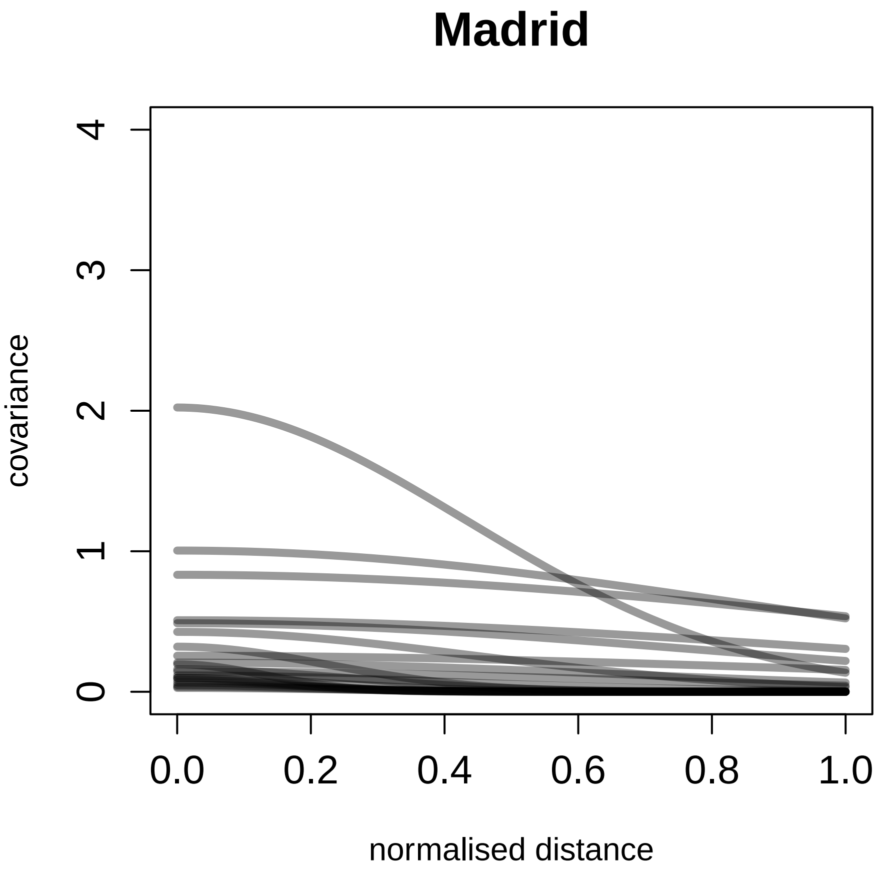


Figure 20: Covariance between parks in Madrid as function of normalised distance..

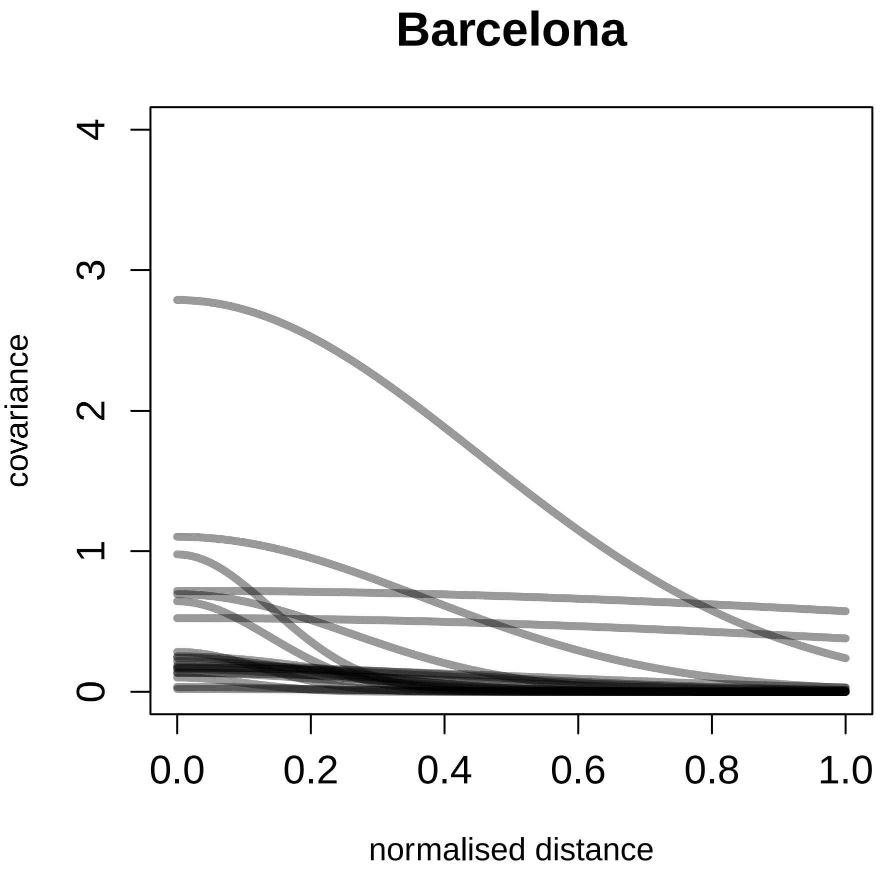


Figure 21: Covariance between parks in Barcelona as function of normalised distance.

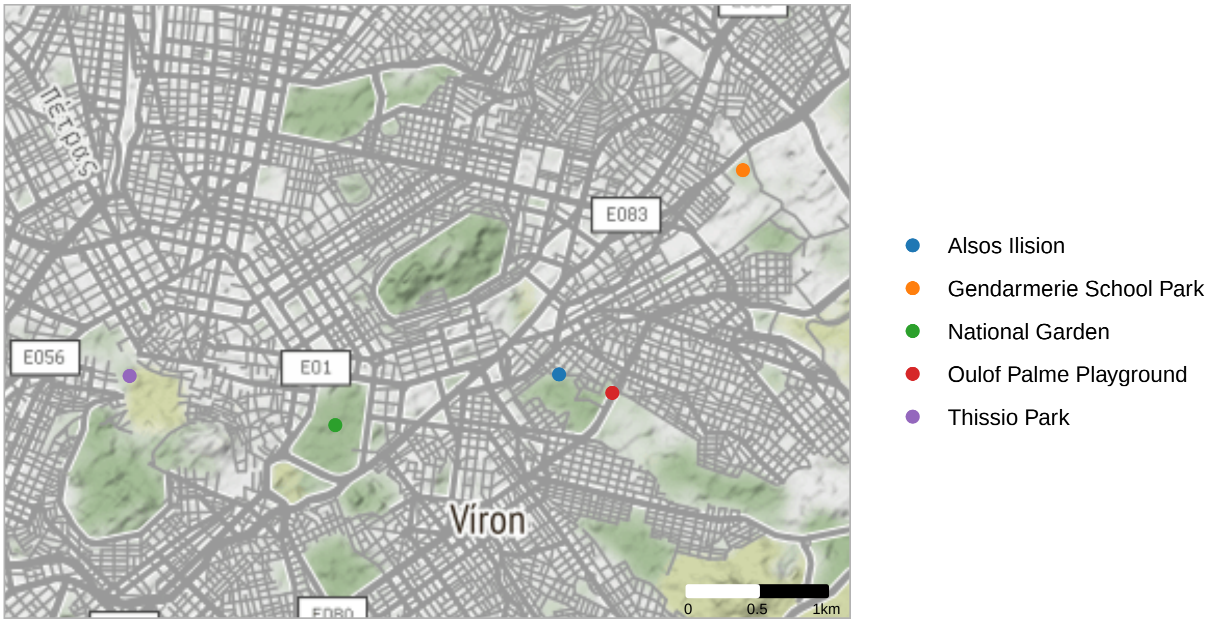


Figure 22: Map of all parks sampled in Athens.

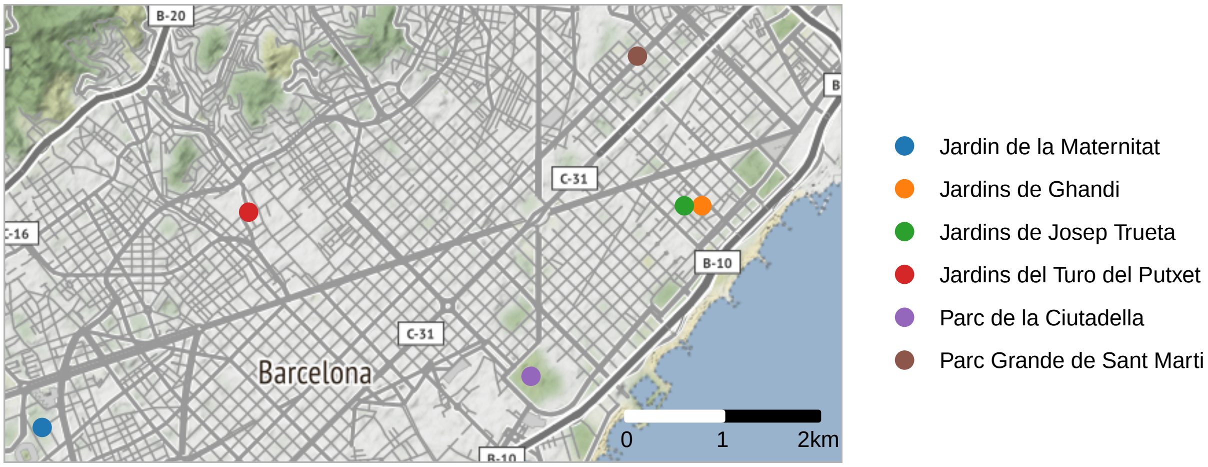


Figure 23: Map of all parks sampled in Barcelona

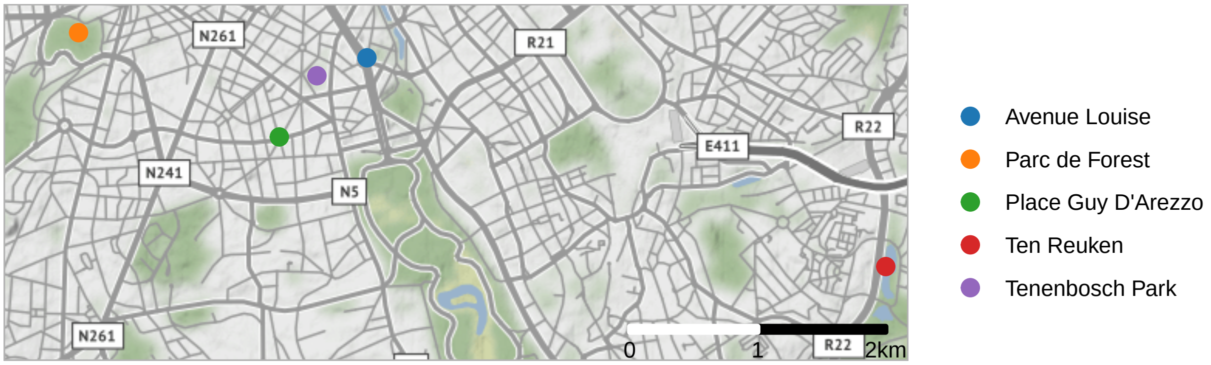


Figure 24: Map of all parks sampled in Brussels

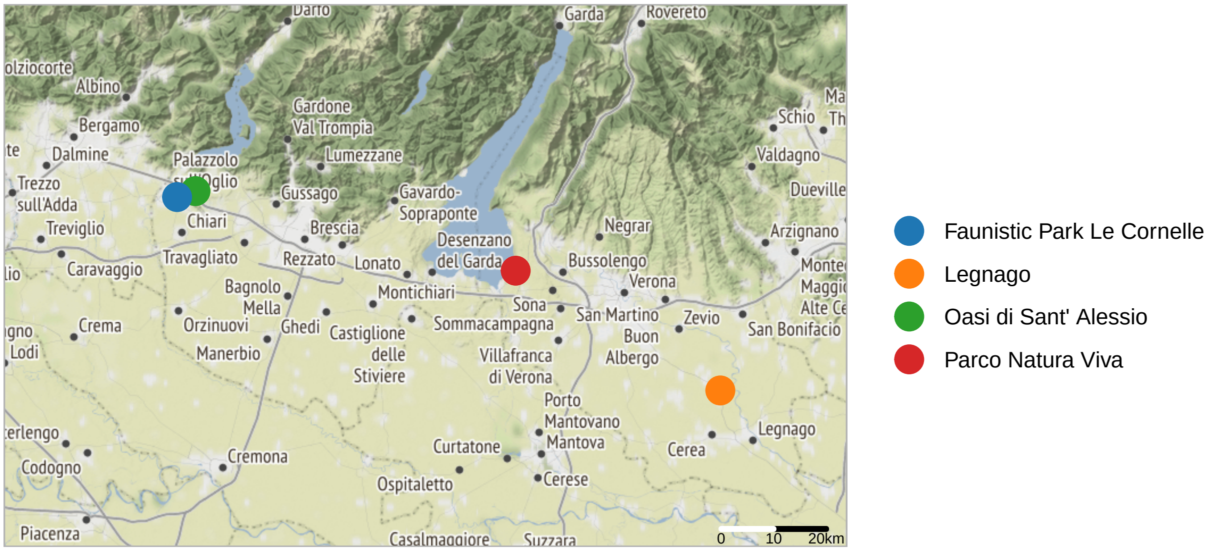


Figure 25: Map of all parks sampled in Italy

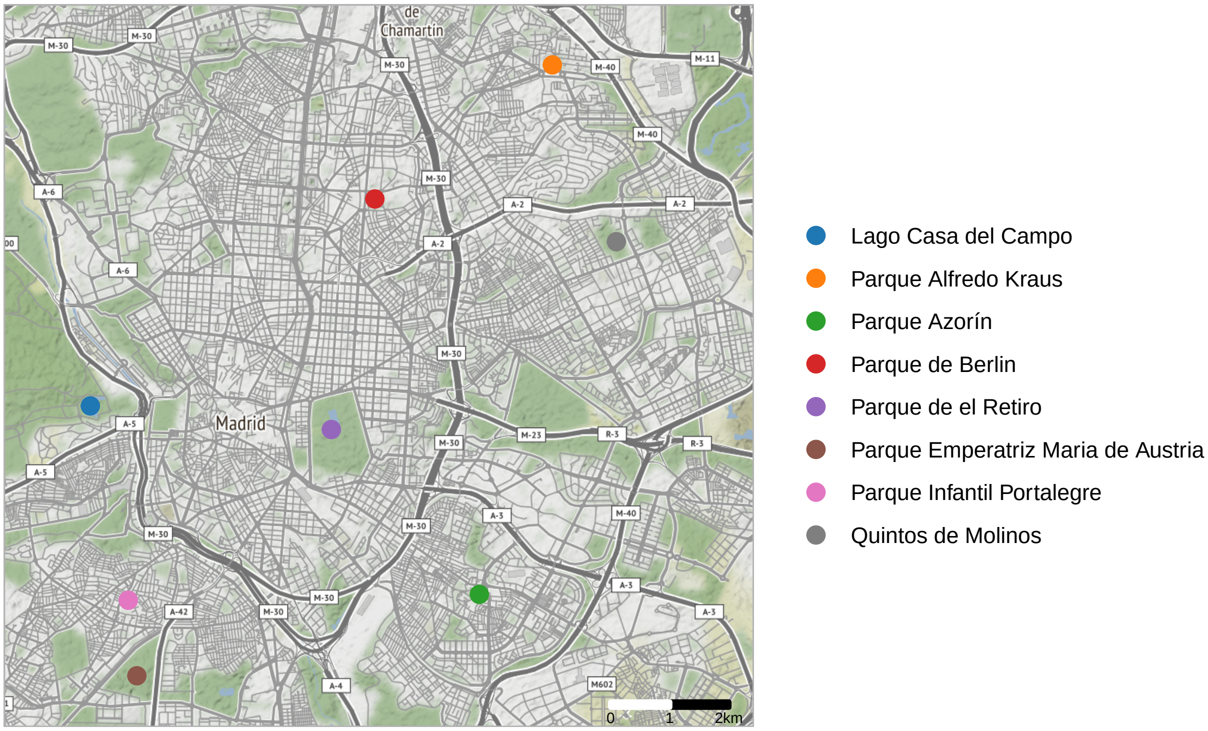


Figure 26: Map of all parks sampled in Madrid
